# Atomistic Profiling of KRAS Interactions with Monobodies and Affimer Proteins Through Ensemble-Based Mutational Scanning Unveils Conserved Residue Networks Linking Cryptic Pockets and Regulating Mechanisms of Binding, Specificity and Allostery

**DOI:** 10.1101/2025.03.11.642708

**Authors:** Mohammed Alshahrani, Vedant Parikh, Brandon Foley, Guang Hu, Gennady Verkhivker

## Abstract

KRAS, a historically “undruggable” oncogenic driver, has eluded targeted therapies due to its lack of accessible binding pockets in its active state. This study investigates the conformational dynamics, binding mechanisms, and allosteric communication networks of KRAS in complexes with monobodies (12D1, 12D5) and affimer proteins (K6, K3, K69) to characterize the binding and allosteric mechanisms and hotspots of KRAS binding. Through molecular dynamics simulations, mutational scanning, binding free energy analysis and network-based analyses, we identified conserved allosteric hotspots that serve as critical nodes for long-range communication in KRAS. Key residues in β-strand 4 (F78, L80, F82), α-helix 3 (I93, H95, Y96), β-strand 5 (V114, N116), and α-helix 5 (Y157, L159, R164) consistently emerged as hotspots across diverse binding partners, forming contiguous networks linking functional regions of KRAS. Notably, β-strand 4 acts as a central hub for propagating conformational changes, while the cryptic allosteric pocket centered around H95/Y96 positions targeted by clinically approved inhibitors was identified as a universal hotspot for both binding and allostery. The study also reveals the interplay between structural rigidity and functional flexibility, where stabilization of one region induces compensatory flexibility in others, reflecting KRAS’s adaptability to perturbations. We found that monobodies stabilize the switch II region of KRAS, disrupting coupling between switch I and II regions and leading to enhanced mobility in switch I of KRAS. Similarly, affimer K3 leverages the α3-helix as a hinge point to amplify its effects on KRAS dynamics. Mutational scanning and binding free energy analysis highlighted the energetic drivers of KRAS interactions. revealing key hotspot residues, including H95 and Y96 in the α3 helix, as major contributors to binding affinity and selectivity. Network analysis identified β-strand 4 as a central hub for propagating conformational changes, linking distant functional sites. The predicted allosteric hotspots strongly aligned with experimental data, validating the robustness of the computational approach. Despite distinct binding interfaces, shared hotspots highlight a conserved allosteric infrastructure, reinforcing their universal importance in KRAS signaling. The results of this study can inform rational design of small-molecule inhibitors that mimic the effects of monobodies and affimer proteins, challenging the “undruggable” reputation of KRAS.

## Introduction

The RAS oncogene family, comprising KRAS, HRAS, and NRAS, plays a critical role in cancer development, with KRAS being the most frequently mutated member. It is implicated in approximately 30% of all RAS-driven cancers, while HRAS and NRAS mutations are less common, occurring in about 8% and 3% of cases, respectively.^1–8^ RAS proteins act as molecular switches, toggling between an active GTP-bound state and an inactive GDP-bound state.^9^ Oncogenic mutations in RAS, particularly at residues G12, G13, and Q61, disrupt GTP hydrolysis, trapping the protein in its active, GTP-bound conformation.^10–17^ This constitutive activation leads to the persistent overactivation of downstream signaling pathways, including the MAP kinase pathway (RAS-RAF-MEK-ERK) and the PI3K pathway (PI3K-AKT-mTOR). These pathways promote uncontrolled cell proliferation, survival, and metabolic changes, driving tumorigenesis.^18^

The interaction between KRAS and RAF1 (CRAF) is essential for MAPK pathway activation, which governs cell growth, differentiation, and survival. RAF1, a serine/threonine kinase, binds KRAS via its RAS-binding domain (RBD) and cysteine-rich domain (CRD). Structural studies, including high-resolution crystal structures, show that the RBD primarily interacts with KRAS’s switch I (SWI, residues 25–40) and switch II (SWII, residues 60–76), regions that undergo conformational shifts during the GTPase cycle, transitioning between inactive and active states.^19,20^ Using paramagnetic relaxation enhancement (PRE) analyses, the structure of a hetero-tetrameric complex of KRAS with RAF1 RBD and CRD was determined.^21^ This study revealed that RBD binding allosterically generates two distinct KRAS dimer interfaces, dynamically equilibrating between a free KRAS-favored interface and one stabilized by the RBD–CRD complex.

Despite its well-established role as a therapeutic target, targeting KRAS has historically been considered “undruggable” due to its high affinity for guanine nucleotides, lack of deep hydrophobic pockets, and structural plasticity. The recent advances in understanding KRAS allostery and the discovery of allosteric binding pockets have reinvigorated efforts to develop effective inhibitors. The canonical view of KRAS as a globular protein with limited druggable sites has shifted with the identification of dynamic conformational changes in KRAS exposing transient pockets that can serve as targets for small molecules.^11,16^ The SWI/SWII-pocket, exists between the SWI and SWII regions of RAS in the region involved in the binding of the nucleotide exchange factor, Son of Sevenless (SOS). Several groups have independently developed active KRA compounds that bind this pocket with varying affinities. ^22–24^ BI-2852 is a groundbreaking non-covalent, pan-KRAS inhibitor that targets a previously elusive allosteric SWI/SWII site on KRAS, binding this pocket with nanomolar affinity.^24^ Unlike covalent KRAS-G12C inhibitors, which specifically target the mutated G12C residue in its inactive state, BI-2852 is mechanistically distinct as it binds to a conserved pocket present in both the active (GTP-bound) and inactive (GDP-bound) conformations of KRAS. BI-2852 exerts its inhibitory effects by dual mechanisms: it blocks SOS-mediated nucleotide exchange, preventing the transition from the inactive to the active state of KRAS, and it disrupts the interaction between KRAS and its downstream effector proteins.^24^ The pan-RAS monobody, JAM20 binds to all RAS variants (KRAS, HRAS, and NRAS) with nanomolar affinity.^25^ Structural studies using NMR and mutation analysis revealed that JAM20 targets SWI/SWII pocket of RAS, a site similarly recognized by small-molecule inhibitors like BI-2852.^25^ Notably, JAM20 directly competes with both the RAF and BI-2852 for binding to RAS, highlighting the importance of the SI/SII pocket as a druggable site and underscoring the potential for synergistic targeting the SI/SII pocket with diverse modalities, such as monobodies and small molecules. A novel allosteric inhibitor RB-456 binds the GDP-bound and GCP-bound conformation of KRAS-G12D by forming interactions with a dynamic allosteric binding pocket within the SWI/SWII region, exhibiting a significant overlap with the bound conformation of BI-2852 inhibitor.^26^ Isothermal titration calorimetry demonstrated that KRB-456 binds potently to KRAS-G12D with the higher affinity than to KRAS wild-type (KRAS-WT), KRAS-G12V and KRAS-G12C.^26^

Another pocket, known as the SWII pocket (SII-P) was identified using a tethered warhead approach that exploits the reactivity of the cysteine in the G12C mutant.^14,15^. The SII-P is a shallow binding site located between the SWII loop (residues A59–Y64), which engages effector proteins, the α2-helix (S65–T74), and the α3-helix (T87–K104). Accessibility to the SII-P depends on conformational changes in the SWII region (residues 60-76), which vary significantly based on the nucleotide-binding state (GTP vs. GDP). The SII-P becomes accessible when KRAS adopts an inactive GDP-bound conformation or when it is stabilized by specific mutations, such as G12C.^14^ Adagrasib (MRTX849) also targets the SII-P, demonstrating clinical efficacy in KRAS G12C-mutant cancers.^17^ While primarily covalent, ARS-1620 and MRTX849 inhibitors also exhibit allosteric properties by binding outside the nucleotide-binding pocket, effectively disrupting KRAS-driven signaling. The A59/T74 allosteric pocket in KRAS is a relatively less explored but emerging site of interest for targeting KRAS-driven cancers. This pocket is distinct from the well-studied SII-P pocket and is located near the α2-helix and β3-β4 loop of KRAS, involving residues A59 and T74. It has been identified as a potential druggable site due to its role in modulating KRAS dynamics and effector interactions.^27^ The H95/Y96 pocket is located near the α3-helix of KRAS and is adjacent to the nucleotide-binding site. This pocket is formed by residues H95 and Y96, which play a role in stabilizing the GTP-bound conformation. Ligand binding to this site can disrupt nucleotide binding and effector interactions.^28,29^ A cryptic pocket near the P-loop which interacts with the γ-phosphate of GTP can become accessible during conformational changes and can be targeted to disrupt nucleotide binding or hydrolysis.^19^ A membrane-proximal pocket is near the C-terminal hypervariable region (HVR) of KRAS and regulates KRAS membrane association, which is critical for signaling activity.^30^

Monobodies—small, synthetic binding proteins engineered to selectively recognize specific protein conformations—have emerged as a promising tool for targeting the oncogenic mutant KRAS(G12D) with high specificity over KRAS-WT and other oncogenic variants. These monobodies represent a novel class of inhibitors that exploit cryptic or conformationally flexible sites on KRAS, offering a unique approach to disrupt KRAS-driven signaling pathways. Crystallographic studies have revealed that these monobodies bind to the SWII pocket which is a shallow groove formed between the SWII and the α3-helix of KRAS. This pocket is highly dynamic and undergoes significant conformational changes depending on the nucleotide-binding state and presence of mutations such as G12D.^31^ Notably, structural studies revealed that the monobodies captured this pocket in its widely open conformation, highlighting the critical role of conformational flexibility in enabling access to cryptic binding sites. By locking KRAS-G12D in an inactive or less active state, these monobodies effectively inhibit downstream signaling pathways, such as the MAPK and PI3K pathways, which are central to tumorigenesis.^31^

The crystal structures of two monobodies, 12D1 and 12D5 in complex with KRAS-G12D^31^ revealed striking similarities to the binding mode of MRTX1133 inhibitor targeting KRAS-G12D where both monobodies and the inhibitor interact with the side chain of KRAS-D12 that may be crucial for achieving high specificity toward the G12D mutation ^32^ Furthermore, deep mutational scanning (DMS) experiments pointed to key residues within KRAS-G12D that are critical for binding and state-specific selectivity, particularly specific interactions with D12 and H95 positions on KRAS.^32^ One of the most remarkable features of these monobodies is their selectivity for KRAS-G12D over KRAS-WT and other oncogenic mutants G12V or G12C. It was initially suggested that his selectivity may arise from their ability to engage key residues that are uniquely positioned in the G12D mutant but the precise mechanisms and complex interplay of thermodynamic contributions yielding the monobody selectivity remains active area of investigation and is addressed in the current study.

The crystal structures of 12D1 and 12D5 also provided additional snapshots of the SWII pocket contributing to the growing body of atomic-level insights into the plastic nature of this site.^33–37^ Collectively, these studies define the SWII-α3 helix pocket as a highly adaptable region capable of accommodating diverse ligands, including both small molecules and protein-based binders. By stabilizing specific conformations of KRAS, monobodies provide a mechanistic framework for designing allosteric inhibitors that trap KRAS in inactive states.

Affimer proteins, a class of engineered scaffold-based biologics, have emerged as powerful tools for targeting challenging proteins like KRAS. Recent studies have identified two distinct druggable pockets on KRAS using affimers: one associated with inhibiting nucleotide exchange and the other with blocking effector binding.^38^ Affimer K6 binds between SWI and SWII of KRAS, regions critical for nucleotide exchange and effector binding. The variable region 1 of K6 interacts directly with these dynamic regions, stabilizing a conformation that mimics the effects of existing small-molecule inhibitors. Competitive nanoBRET assays confirm that affimer K6 functionally mimics small-molecule inhibitors and disrupts the equilibrium between active (GTP-bound) and inactive (GDP-bound) states of KRAS, effectively trapping it in an inactive state.^38^ Affimer K3 targets a novel druggable pocket located between the SWII region and the α3 helix of KRAS. This pocket, termed the SII/α3 pocket, represents a previously unseen conformer of the SII pocket in KRAS-WT. The SII/α3 pocket coincides with a similar conformer targeted by the cyclic peptide KRpep-2d in the KRAS(G12D) mutant,^39,40^ suggesting that this pocket is conserved across KRAS variants and may be universally applicable for targeting oncogenic mutations. A comprehensive analysis of KRAS energetics and allostery was undertaken using a combination of structural biology and biophysical techniques to map the energetic and allosteric landscape of KRAS.^41^ This seminal study characterized allosteric energy landscapes that highlighted the thermodynamic consequences of mutations across KRAS providing a quantitative framework to determine allosteric binding sites and their effect on KRAS binding.

Computational studies, including molecular dynamics (MD) simulations and free energy analysis, have provided important insights into the structural dynamics, mutational effects, and inhibitor binding mechanisms of KRAS with small molecule inhibitors and effector proteins. MD simulations revealed that oncogenic mutations like Q61H can modulate the flexibility of inactive KRAS without significantly affecting active KRAS form, while Q61A, Q61H, and Q61L modifications increase flexibility of switch regions, disrupting KRAS activity.^12^ Multiscale simulations highlighted how membrane orientation influences KRAS dynamics within the RAS-RBD/CRD complex, with the membrane playing a key role in stabilizing metastable states through interactions between CRAF and KRAS.^42^ MD simulations also explored the dynamics and stability of KRAS-WT and its oncogenic variants (G12C, G12D, G12V, G13D) followed by the free energy landscape analysis revealing unique conformational dynamics and altered thermodynamic stability in the oncogenic KRAS variants.^43^ Multiple MD simulations were performed on KRAS-WT and KRAS mutant structures to investigate how G12C and G12D mutations stabilize the active state and how AMG-510 (Sotorasib) and MRTX1133 inhibitors force these mutants into the inactive state.^45^ The study revealed that binding of Sotorasib can induce significant stabilization of the SWII region of KRAS-G12C, surpassing that of MRTX1133 bound with KRAS-G12D mutant. Microsecond MD simulations and Markov State Model (MSM) analysis probed kinetics of MRTX1133 binding to KRAS G12D revealing the kinetically metastable states and pathways of MRTX1133 binding.^37^ Gaussian accelerated molecular dynamics (GaMD) simulations of the GDP-bound G12A, G12D, and G12R KRAS variants to probe mutation-mediated impacts on conformational alterations of KRAS, showing that all three G12 mutations can alter the structural flexibility of the switch domains.^45^

Despite a significant body of structural, biochemical and computational studies of KRAS dynamics and binding there are a number of open questions concerning the mechanistic aspects of KRAS binding and allosteric interactions with monobodies and affimer proteins. Among questions addressed in our current study are the following: How do monobodies and affimers induce allosteric changes in KRAS and what are the dynamic effects of allosteric modulation on KRAS function? What is the mechanism and key binding hotspots and interactions that drive selectivity of monobodiesAre there conserved allosteric networks across KRAS mutants? What are the key determinants of high-affinity binding between KRAS and monobodies and affimers and which KRAS sites can serve as hotspots for both binding and allostery?

In this study, we explore some these issues in detail by employing an integrative computational simulation strategy that combined microsecond MD simulations, binding free energy analysis and network modeling of allostery for KRAS complexes with a panel of monobodies and affimer proteins targeting distinct allosteric pockets on KRAS. The detailed MM-GBSA analysis of binding energetics reveals the thermodynamic drivers of binding and the energetic contributions of the binding affinity hotspots to KRAS stability and effector binding, showing agreement with the experimental data. Using MD-inferred conformational ensembles and dynamics-based network modeling, this study mapped potential allosteric hotspots and allosteric communication pathways in KRAS complexes. Consistent with rigorous experimental analysis, our results confirmed that the central β-strands of KRAS acts as a hub for transmitting allosteric signal between distant functional sites, regulating allosteric communication between district allosteric pockets of KRAS and binding partners. The detected maps of allosteric sites and binding interface hotspots in the KRAS complexes highlight role of conserved hotspots binding and allostery that enable diverse allosteric pathways and mechanisms of targeting allosteric pockets.

## Materials and Methods

### Structural Analysis and All-Atom MD Simulations of KRAS Complexes with Monobodies and Affimer Proteins

The crystal and cryo-EM structures of the KRAS-G12D complexes with monobody 12D1 (pdb id 8EZG) and monobody 12D5 (pdb id 8F0M) as well as KRAS complexes with affimer K6 (pdb id 6YR8), affimer K3 (pb id 6YXW) and affimer K69 (pdb id 7NY8) are obtained from the Protein Data Bank.^46^ For simulated structures, hydrogen atoms and missing residues were initially added and assigned according to the WHATIF program web interface.^47^ Missing residues or loops were modeled using MODELLER^48^, and the structures were cleaned to remove water molecules, ligands, and non-standard residues. The tleap module in the AMBER package was also employed to add the missing hydrogen atoms of the structures.^49^ The missing regions are subsequently reconstructed and optimized using template-based loop prediction approach ArchPRED.^50^ The side chain rotamers were refined and optimized by SCWRL4 tool.^51^ Protonation states of ionizable residues were assigned using PROPKA, ensuring histidine residues were in the appropriate tautomeric state.^52,53^ The tleap module in AMBER was used to generate the topology (prmtop) and coordinate (inpcrd) files. The system was solvated in a TIP3P water box with a 12 Å buffer and neutralized using Na+ or Cl-ions. Molecular mechanics parameters of proteins were assigned according to the ff14SB force field.^54–56^ Two stages of minimization were performed: (1) with restraints on the protein backbone to relax solvent and ions, and (2) without restraints to minimize the entire system. The minimization was conducted using the pmemd.cuda module with the following parameters: 1,000 cycles of steepest descent followed by 500 cycles of conjugate gradient minimization, a cutoff of 10 Å for non-bonded interactions, and the ff14SB force field for proteins.^54,55^ The relaxation process includes minimization, heating, restriction run, and equilibrium run. After minimization, the system is heated from 100K to 300K, over 1 nanosecond of simulation time at a constant volume, with integration time 1 fs. Then we relax system at a constant pressure with protein restraints over 1 ns of simulation time at a constant pressure with restraints set to 1 kcal/mol Å^2. Finally, we relax the system with no restraints for 1 ns of simulation time at a constant pressure. Production MD simulations were performed for 2µs under NPT conditions (300 K, 1 bar) using the pmemd.cuda module. The temperature was maintained using the Langevin thermostat, and pressure was controlled with the Monte Carlo barostat. A time step of 2 fs was used, and bonds involving hydrogen atoms were constrained using the SHAKE algorithm. MD trajectories were saved every 100 ps for analysis. The ff14SB force field was used for proteins, and the TIP3P model was used for water. For each system, MD simulations were conducted three times in parallel to obtain comprehensive sampling. Each individual simulation has 10,000 frames. The CPPTRAJ software in AMBER 18 was used to calculate the root mean squared fluctuation (RMSF) of MD simulation trajectories in which the initial structure was used as reference.^57,58^ The structures of were visualized using Visual Molecular Dynamics (VMD 1.9.3) ^59^ and PyMOL (Schrodinger, LLC. 2010. The PyMOL Molecular Graphics System, Version X.X.)

### Mutational Scanning Analysis of Binding Interfaces in the KRAS Protein Complexes

We conducted a systematic mutational scanning analysis of the KRAS residues in the KRAS complexes using conformational ensembles of KRAS complexes and averaging of mutation-induced energy changes. Every KRAS residue was systematically mutated using all substitutions and corresponding protein stability and binding free energy changes were computed with the knowledge-based BeAtMuSiC approach.^60,61^ This approach is based on statistical potentials describing the pairwise inter-residue distances, backbone torsion angles and solvent accessibilities, and considers the effect of the mutation on the strength of the interactions at the interface and on the overall stability of the complex. The binding free energy of protein-protein complex can be expressed as the difference in the folding free energy of the complex and folding free energies of the two protein binding partners:

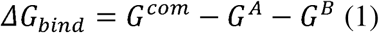

The change of the binding energy due to a mutation was calculated then as the following:

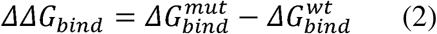

We leveraged rapid calculations based on statistical potentials to compute the ensemble-averaged binding free energy changes using equilibrium samples from simulation trajectories. The binding free energy changes were obtained by averaging over 1,000 equilibrium samples for each of the systems studied.

### MM-GBSA Binding Free Energy Computations of the KRAS Complexes with Monobodies and Affimer Proteins

We calculated the ensemble-averaged changes in binding free energy using 1,000 equilibrium samples obtained from simulation trajectories for each system under study. The binding free energy of the KRAS complexes were assessed using the MM-GBSA approach.^62,63^ The energy decomposition analysis evaluates the contribution of each amino acid to binding of KRAS to proteins.^64,65^ The binding free energy for the KRAS complex was obtained using:

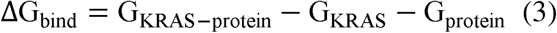

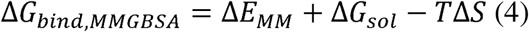

where Δ*E_MM_*is total gas phase energy (sum of Δ*E_internal_*, Δ*E_electrostatic_*, and Δ*Evdw*); Δ*Gsol* is sum of polar (Δ*G_GB_*) and non-polar (Δ*G_SA_*) contributions to solvation. Here, G_KRAS–protein_ represent the average over the snapshots of a single trajectory of the complex, G_KRAS_ and G_prorein_ corresponds to the free energy of KRAS binding with monobodies (12D1, 12D5) and affimer proteins (K6, K3, K69). The polar and non-polar contributions to the solvation free energy is calculated using a Generalized Born solvent model and consideration of the solvent accessible surface area.^66^ MM-GBSA is employed to predict the binding free energy and decompose the free energy contributions to the binding free energy of a protein–protein complex on per-residue basis. The binding free energy with MM-GBSA was computed by averaging the results of computations over 1,000 samples from the equilibrium ensembles. We employ the “single-trajectory” (one trajectory of the complex) protocol for the computations. To reduce the noise in calculations, it is common that each term is evaluated on frames from a single MD trajectory of the bound complex. In this study, we choose the “single-trajectory” protocol, because it is less noisy due to the cancellation of intermolecular energy contributions. This protocol applies to cases where significant structural changes upon binding are not expected. Entropy calculations typically dominate the computational cost of the MM-GBSA estimates. Therefore, it may be calculated only for a subset of the snapshots, or this term can be omitted.^67,68^ However, for the absolute affinities, the entropy term is needed, owing to the loss of translational and rotational freedom when a complex with a binding partner is formed. In this study, the entropy contribution was not included in the calculations of binding free energies of the complexes because the entropic differences in estimates of relative binding affinities are expected to be small owing to small mutational changes and preservation of the conformational dynamics.^67,68^ MM-GBSA energies were evaluated with the MMPBSA.py script in the AmberTools21 package.^69^

### Graph-Based Dynamic Network Analysis of Protein Ensembles

To analyze protein structures, we employed a graph-based representation where residues are modeled as network nodes, and non-covalent interactions between residue side-chains define the edges. This approach captures the spatial and functional relationships between residues, providing insights into the protein’s structural and dynamic properties. The graph-based framework allows for the integration of both structural and evolutionary information, enabling a comprehensive analysis of residue interactions. The residue interaction networks were constructed by defining edges based on non-covalent interactions between residue side-chains.^70,71^ The weights of these edges were determined using two key metrics: (a) dynamic residue cross-correlations derived from MD simulations where these correlations quantify the coordinated motions of residue pairs^72^ and (b) coevolutionary couplings measured using mutual information scores, where these couplings reflect evolutionary constraints and residue-residue dependencies.^73^ The edge lengths (weights) between nodes ii and jj were computed using generalized correlation coefficients, which integrate both dynamic correlations and coevolutionary mutual information. Only residue pairs observed in at least one independent simulation were included in the network. The matrix of communication distances was constructed using generalized correlations between composite variables that describe both the dynamic positions of residues and their coevolutionary mutual information. This matrix provides a quantitative measure of communication efficiency between residues, reflecting both their physical proximity and evolutionary relationships. The Residue Interaction Network Generator (RING) program^74–77^ was used to generate residue interaction networks from the conformational ensemble. The edges in these networks were weighted to reflect the frequency of interactions observed in the ensemble. Network files in XML format were generated for all structures using the RING v3.0 webserver. Network graph calculations were performed using the Python package NetworkX.^78,79^ This included the computation of key network parameters, such as shortest paths and betweenness centrality, to identify residues critical for communication within the protein structure. The short path betweenness (SPC) of residue *i* is defined to be the sum of the fraction of shortest paths between all pairs of residues that pass through residue *i*:

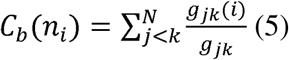

where *g_jk_*denotes the number of shortest geodesics paths connecting *j* and *k*, and *g _jk_* (*i*) is the number of shortest paths between residues *j* and *k* passing through the node *n*_i_. Residues with high occurrence in the shortest paths connecting all residue pairs have a higher betweenness values. For each node *n*, the betweenness value is normalized by the number of node pairs excluding *n* given as (*N* −1)(*N* - 2) / 2, where *N* is the total number of nodes in the connected component that node *n* belongs to. To account for differences in network size, the betweenness centrality of each residue ii was normalized by the number of node pairs excluding ii. The normalized short path betweenness of residue *i* can be expressed as:

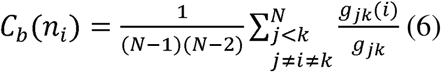

*g_jk_* is the number of shortest paths between residues *j* and k; *g _jk_* (*i*) is the fraction of these shortest paths that pass through residue *i*.

Residues with high normalized betweenness centrality values were identified as key mediators of communication within the protein structure network. These residues are likely to play critical roles in maintaining the protein’s structural integrity and functional dynamics.

### Network-Based Mutational Profiling of Allosteric Residue Centrality

Through mutation-based perturbations of protein residues we compute dynamic couplings of residues and changes in the short path betweenness centrality (SPC) averaged over all possible modifications in a given position.

The change of SPC upon mutational changes of each node is reminiscent to the calculation of residue centralities by systematically removing nodes from the network.

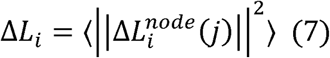

where *i* is a given site, *j* is a mutation and 〈⋯〉denotes averaging over mutations. 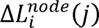 describes the change of SPC parameters upon mutation U in a residue node i. ΔL_i_ is the average change of ASPL triggered by mutational changes in position i.

Z-score is then calculated for each node as follows:

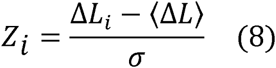

(ΔL) is the change of the SPC network parameter under mutational scanning averaged over all protein residues and σ is the corresponding standard deviation. The ensemble-average Z score changes are computed from network analysis of the conformational ensembles of KRAS-RAF1 complexes using 1,000 snapshots of the simulation trajectory. Through this approach, we evaluate the effect of mutations in the KRAS residues on allosteric communications with binding proteins.

## Results

### Conformational Dynamics and Cooperative Motions of the KRAS Complexes with Monobodies

A complete GTPase reaction requires well-ordered conformations of the protein active site, which includes the phosphate-binding loop, P-loop (residues 10–17), switch I (residues 25–40) and switch II (residues 60–76) regions (Figure 1). The major elements of KRAS include the P-loop (residues 10-17), Switch I (residues 25-40) and Switch II regions (residue 60-76), the α-helices (α-helix 1: 15–24; α-helix 2: 67–73; α-helix 3: 87–104; α-helix 4: 127–136; α-helix 5:

**Figure 1.**
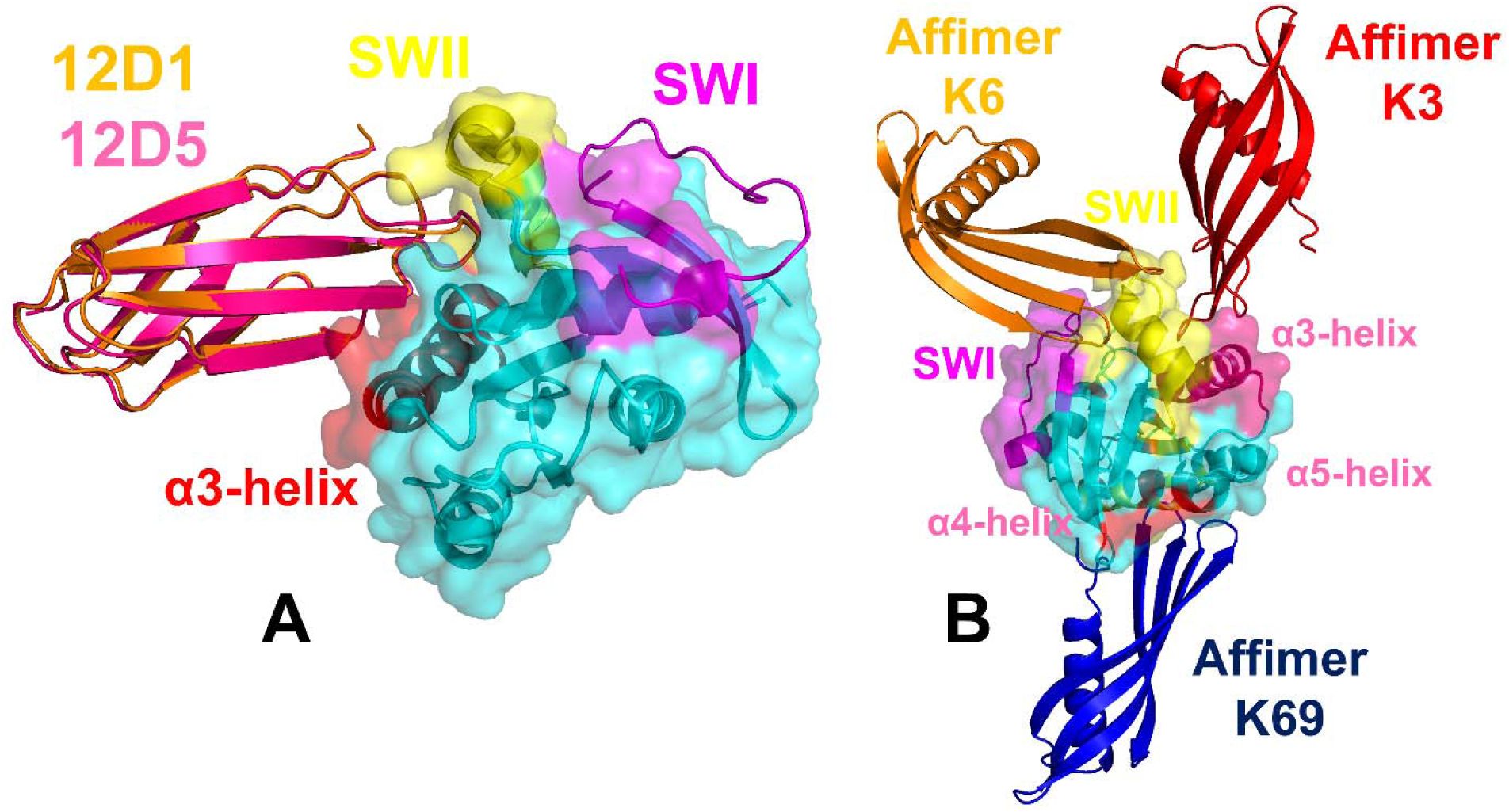
Structural overview and organization of the KRAS protein complexes with monobodies 12D1/12D5 and affimer proteins K6, K3 and K69. (A) Structural overlay of the crystal structures of KRAS-G12D protein with monobodies 12D1 (pdb id 8EZG) and 12D5 (pdb id 8F0M). KRAS protein is shown in cyan ribbons and transparent surface. The highlighted functional regions of KRAS are annotated: switch I, SW1 (residues 24-40 in magenta ribbons), switch II, SWII (residues 60-76 in yellow ribbons), allosteric KRAS lobe (residues 87-166 in yellow ribbons), the α-helix 3 (residues 87-104 in red ribbons). The monobody 121 is in orange ribbons and monobody 12D5 is in hotpink-colored ribbons. (B) Structural overlay of the crystal structures of KRAS protein with affimer proteins K6 (pdb id 6YR8), K3 (pdb id 6YXW) and K69 (pdb id 7NY8). KRAS proteins are shown in cyan ribbons and transparent surface. The highlighted functional regions of KRAS are annotated: switch I, SW1 (residues 24-40 in magenta ribbons), switch II, SWII (residues 60-76 in yellow ribbons), allosteric KRAS lobe (residues 87-166 in yellow ribbons), the α-helix 3 (residues 87-104 in red ribbons), α-helix 4 (residues 127–136 and α-helix 5 (residues 148–166) are in pink-colored ribbons. Affimer K6 is shown in orange ribbons, affimer K3 is in red ribbons and affimer K69 is in blue ribbons. Affimer K6 binds to a shallow hydrophobic pocket on KRAS between the SWI and SWII regions. Affimer K3 binds to a pocket between the SWII and α3 helix. Affimer K69 (in blue) binds the allosteric lobe between helices 4 and 5 on the opposite side of KRAS (in cyan) to affimers K3 (in red) and K6 (in orange).

148–166) and β-strands (β-strand 1: 3–9; β-strand 2: 38–44; (β-strand 3: 51–57; β-strand 4: 77–84; β-strand 5: 109–115; β-strand 6: 139–143) and the allosteric lobe (residues 87-166).

The conformational flexibility of the KRAS was analyzed by calculating the root mean square fluctuations (RMSF) distribution for the KRAS residues in different complexes (Figure 2A,B). We also characterized essential motions and determined the hinge regions in the KRAS complexes using principal component analysis (PCA) of trajectories using the CARMA package.^80^ The local minima along these profiles are typically aligned with the immobilized in global motions hinge centers, while the maxima correspond to the moving regions undergoing concerted movements leading to global changes in structure.^81,82^ The low-frequency ‘soft modes’ are characterized by their cooperativity and there is a strong relationship between conformational changes and the ‘soft’ modes of motions intrinsically accessible to protein structures.^81,82^

**Figure 2.**
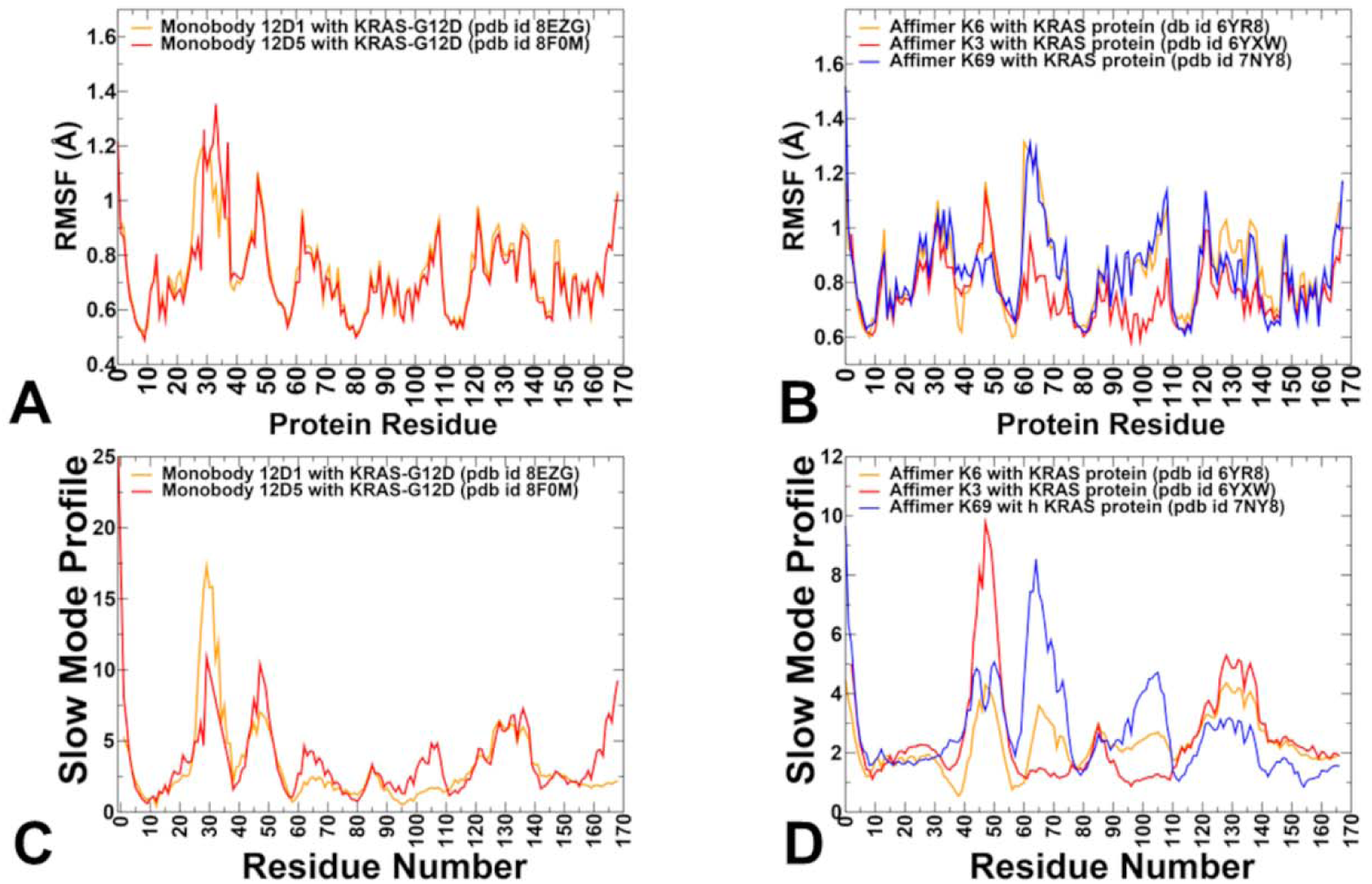
Conformational dynamics profiles and essential mobility profiles of the KRAS protein complexes. (A) The RMSF profiles of KRAS-G12D protein residues obtained from MD simulations of the KRAS complexes with monobodies 12D, pdb id 8EZG (in orange lines) and 12D5, pdb id 80M (in red lines). (B) The RMSF profiles of KRAS protein residues obtained from MD simulations of the KRAS complexes with affimer protein K6, pdb id 6YR8 (in orange lines), affimer K3, pdb id 6YXW (in red lines) and affimer K69, pdb id 7NY8 (in blue lines). (C) Functional dynamics of the KRAS-G12D protein in complexes is presented as the essential mobility profiles with monobodies 12D1 (in orange) and 12D (in red). The essential mobility profiles are averaged over the first three major low frequency modes. (D) The essential mobility/slow mode profiles of KRAS in complexes with affimer proteins K6 (in orange), K3 (in re) and K69 (in blue).

The crystal structure shows that monobody 12D1 binds primarily to the cleft formed between the SWII region and the α3 helix of KRAS-G12D (Figure 1A). This pocket exhibits unique conformations and interactions due to the G12D mutation. While other KRAS-G12D specific inhibitors (e.g., KD2 peptide, KRpep-2d and MRTX1133) also target the SWII pocket, 12D1 occupies a larger volume within the SWII pocket, displacing the SWII region closer toward the SWI (Figure 1A). Monobody 12D5 targets the H95/Y96 allosteric pocket, located near the α3 helix (residues 87–104) and adjacent to the Switch regions (Figure 1A).

The RMSF profiles for the KRAS complexes with monobodies 12D1 and 12D5 showed considerable similarity revealing dynamic signatures sable over the simulation timescale, where SWI (residues 25-40) displayed greater mobility compared to other regions (Figure 2A). Interestingly, and SWII region (residues 60-76) showed greater stability, particularly around residues 70-76 (Figure 2A). Conformational dynamics profiles also highlighted minor thermal fluctuations and stability of the α-helix 3 (residues 87–104). The P-loop is a highly conserved region in KRAS that interacts with the phosphate groups of GTP. It plays a critical role in nucleotide binding and stabilization. MD simulations showed that the P-loop (residues 10-17) exhibits moderate flexibility that allows the P-loop to accommodate conformational changes (Figure 2A). MD simulations also revealed that 12D5 stabilizes the α3 helix (residues 87-104), reducing its flexibility and altering the local environment of the H95/Y96 pocket. Perturbations at this site propagate through β-strands 2 and 3 (residues 38–44 and 51–57), which are coupled to the α3 helix (Figure 2A).

PCA analysis of slow mode profiles for KRAS binding with monobodies demonstrated an interesting compensatory behavior of SWI and SWII movements where binding of monobody 12D1 can cause the greater immobilization of the SWII region that enables larger functional displacements of the SWI (Figure 2B). This contrasts to the slow mode profile of KRAS binding with monobody 12D5 that can effectively suppress functional movements of SWI and allow only moderate fluctuations of thw SWII region (Figure 2B). The differential behavior of SWI and SWII in the KRAS complexes with monobodies 12D1 and 12D5 revealed by MD simulations and PCA—where SWI exhibits enhanced mobility in the complex with 12D1 while SWII becomes more rigid —can be explained by the combination of structural, allosteric, and functional factors. SWI is an inherently flexible region that undergoes significant conformational changes depending on the nucleotide-binding state (GTP vs. GDP) and interactions with regulatory proteins. In the GTP-bound active state, SWI adopts a well-defined conformation that facilitates effector binding. However, in the GDP-bound inactive state or when bound to monobody 12D1, SWI becomes more dynamic (Figure 2A). The SWI residues 25–40 are directly involved in forming the effector-binding interface and interact closely with the nucleotide-binding pocket, making them sensitive to perturbations in KRAS dynamics induced by binding with monobodies While SWII is another dynamic region of KRAS its mobility is reduced upon binding to both 12D1 and 12D5. This stabilization occurs because monobodies directly interact with the SWII pocket. By rigidifying SWII, monobodies 12D1/12D5 can prevent KRAS from adopting conformations required for effector binding or nucleotide exchange. The rigidification of SWII disrupts the normal coupling between SWI and SWII, leading to compensatory flexibility in SWI in binding with 12D1. As a result, residues 25–40 in SWI can explore a broader range of conformations, resulting in enhanced mobility in slow modes for the KRAS complex with monody 12D1 (Figure 2A,B). The increased mobility of SWI residues 25–40 likely reflects a compensatory mechanism to maintain some degree of functional adaptability despite the immobilization of SWII. This aligns with the inhibitory role of monobody 12D1, which prevents KRAS from engaging with downstream signaling pathways.

Despite similar binding modes and conformational dynamics profiles, the distribution of functional motions can djust in the KRAS complex with monobody 12D5 (Figure 2B). The stabilization of the H95/Y96 pocket and targeting of α3-helix (87–104) can introduce mechanical stress into the KRAS structure, which is redistributed through coordinated motions of β-strand 2: 38–44 and β-strand 3: 51–57 (Figure 2B). This redistribution is evident in the slow mode profiles, where both SWI region and residues 38–60 dominate the slow modes due to their collective movements in the KRAS complex with monobody 12D5 (Figure 2B). At the same time, we observed greater stabilization of the SWII region in the KRAS complex with 12D5. This coupling ensures that perturbations in one region are transmitted to others, enabling a coordinated response to ligand binding. To conclude, our results suggest that monobody 12D1 stabilizes SWII, disrupting the coupling between SWI and SWII regions and leading to compensatory flexibility in SWI. Binding of monobody 12D5 can induce stabilization of the SWII and α3 helix regions and indirectly modulate functional movements of SWI through long-range communication enabled by functional adjustments of β-strands 2 and 3 (Figure 2B). The collective movements of β-strands 2 and 3 may facilitate the adoption of specific conformations for SWI and SWII in the complex with 12D5.

Structural mapping of slow modes on the crystal structures of the KRAS complexes with 12D1 and 12D5 highlighted similarities but also pointed to the overall increased adaptability of KRAS in the complex with 12D5, particularly in the SWI and SWII regions as well as β-strands 2 and 3 (Supporting Information, Figure S1). This is consistent with experimental structural data showing that KRAS structures are nearly identical in complexes with 12D1 and 12D5 except for the SWI region as 12D5 can bind to both nucleotide states of KRAS-G12D.^31^ These delicate modulations of cooperative motions can reflect the allosteric nature of binding mechanisms with monobody proteins that can uniquely affect allosteric signals between different functional regions of KRAS and inhibiting KRAS functions.

### Distinct Dynamic Signatures of KRAS Binding to Affimer Proteins

We also examined conformational dynamics of KRAS binding to affimers K6, K3, and K69. These affimers target distinct regions of KRAS, inducing unique conformational changes that modulate its activity (Figure 1B). The crystal structure of affimer K6 in complex with GDP-bound KRAS-WT was determined at 1.9 Å resolution, revealing that affimer K6 binds to the SI/SII hydrophobic pocket on KRAS (Figure 1B).^38^ The SWI/SWII hydrophobic pocket is formed by residues from both Switch I and Switch II, such as L6, V7, T35, E37, D38, S39, Y40, V44, D54, I55, L56, M67, Y71, T74, and G75.^38^ Structural studies showed that affimer K6 can form a hydrophobic cluster and primarily bind to I36, L56, M67, and Y71 sites on KRAS from SWI and SWII regions. MD simulations of the KRAS-K6 complex revealed that affimer K6 can have larger stabilizing effect on SWI as most of the SWI/SWII hydrophobic pocket positions include SWI residues and yet enable only moderate thermal mobility of SWII inactive conformation (Figure 2C). Affimer K3 binds to the cryptic SWII/α3 pocket, a region that includes parts of the SII region and the α3-helix (residues 87–104).^38^ The RMSF profile for KRAS bound to affimer K3 shows the largest degree of stabilization among the three affimers (Figure 2C). In particular, binding of K3 rigidifies α-helix 2 (residues 67–73) and α-helix 3 (residues 87–104), reducing their flexibility and stabilizing the interaction with the affimer. Unlike affimers K6 and K3, K69 does not stabilize cryptic pockets but instead targets a distinct allosteric site on KRAS.^38^ Conformational dynamics profiles suggested that affimer K69 can induce local and global conformational changes in KRAS, affecting multiple regions simultaneously. The RMSF profiles showed that binding of K69 stabilizes α-helices 4 (residues 127–136) and 5 (residues 148–166) but surprisingly allows for more plasticity in these regions as compared to affimer K3 that binds in a different site (Figure 2C).

PCA identifies slow modes associated with binding of affimer proteins revealing considerable differences in modulation of functional movements induced by K3, K6 and K69 affimers (Figure 2D). For the KRAS complex with K6 the dominant slow modes reflect restricted movements in the SWI and SWII regions due to binding to the SWI/SWII hydrophobic pocket, preventing transitions to active states. Structural mapping of the slow modes averaged over the slowest three modes projected onto the crystal structures of the KRAS complexes with affimer proteins illustrated these observations (Supporting Information, Figure S2). Overall, the analysis of conformational dynamics and slow mode profiles suggested the direct stabilization of the switch regions by affimer K6 may lock KRAS in an inactive conformation and prevent KRAS from adopting conformations required for effector binding or nucleotide exchange.

Binding of K3 rigidifies the SWII region and α3 helix, considerably reducing their flexibility in slow modes. This rigidification acts as a hinge point for transmitting conformational changes across KRAS and can also restrict movements of the SWI regions (Figure 2D). These results are consistent with a similar effect of monobody 12D5, suggesting targeting the cryptic SWII/α3 pocket may disrupt normal coupling between functional regions, leading to enhanced functional displacements β-strands 2 and 3 (residues 38-60) serving as transmitters of allosteric changes induced by affimer K3. This mechanism highlights the importance of the SII/α3 pocket as a target for modulating KRAS dynamics. Indeed, binding of K3 rigidifies KRAS in a conformation harboring a SII/α3 pocket and shows a similar in vitro potency to AMG-510.^38^ The experimental functional studies suggested that targeting this region is a strong approach for allosteric inhibition of KRAS. ^38^ Our analysis of conformational and functional dynamics indicated that SII/α3 pocket is allosterically coupled with movements β-strands 2 and 3 which collectively control allosteric changes and interactions in the KRAS.

The analysis of essential mobility profiles in the KRAS complex with affimer K69 that binds to the allosteric lobe between helices 4 and 5 showed an interesting and unique pattern of collective motions (Figure 2D). Collective displacements and functional motions are observed in SWII region (residues 60-76) and α-helix 3 (residues 87–104), which is “released” from its immobilized conformation seen in the complex with K3. The stabilization of helices 4 and 5 by affimer K69 redistributes conformational sampling to β-sheets 2 and 3. Hence, binding of K69 may lead to more global conformational redistributions transmitted from the allosteric lobe to distal regions (e.g., β-sheets 2 and 3, SWI/SWII), influencing overall protein function. The long-range perturbation of SWI and SWII dynamics induced by affirmer K69 can potentially impair the formation of the effector-binding interface, preventing KRAS from interacting with downstream effectors. Overall, the results showed that the conformational dynamics induced by Affimers K6, K3, and K69 reflect the versatility of KRAS as a target for allosteric modulation. Each affimer employs a distinct mechanism to stabilize inactive states.

### Mutational Profiling of KRAS Binding Interactions with Monobodies

Using conformational ensembles obtained from MD simulations, we performed systematic mutational scanning of the KRAS residues in the KRAS complexes with monobodies 12D1 and 12D5 (Figure 3). Computational mutational scanning was done by averaging the binding free energy changes over the equilibrium ensembles and allows for predictions of the mutation-induced changes of the binding interactions and the stability of the complex. To provide a systematic comparison, we constructed mutational heatmaps for the KRAS interface residues. We used BeatMusic^60,61^ and PRODIGY contact predictor^83,84^ to identify binding interface residues. Residues are considered part of the interface if they are within a defined cutoff distance (typically 5 Å) from atoms in the binding partner.

**Figure 3.**
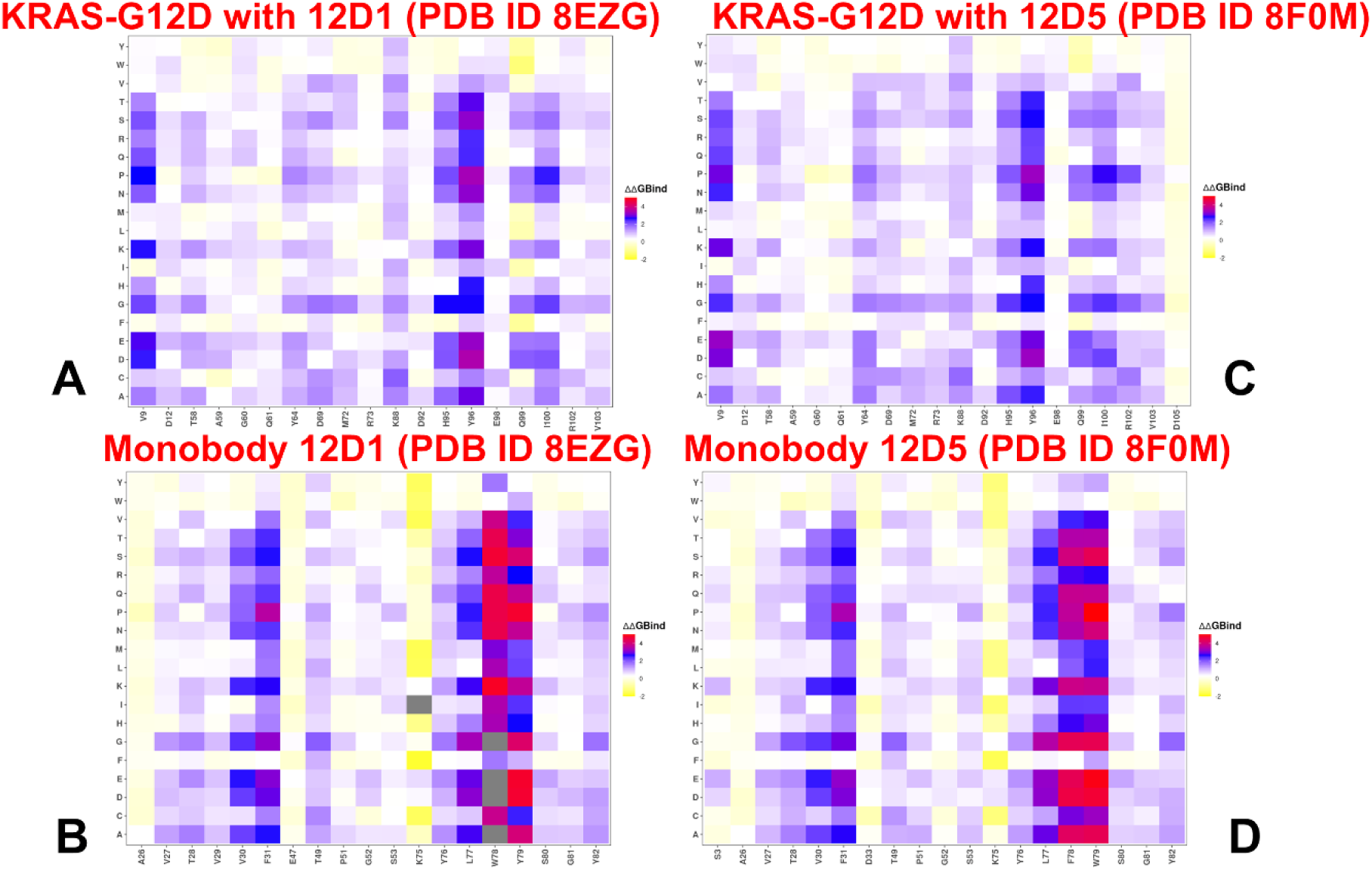
The ensemble-based mutational scanning of binding for the KRAS-G12D complexes with monobodies 12D1 and 12D5. The mutational scanning heatmap for the KRAS binding epitope residues in the complex with 12D1 (A) and heatmap for 12D1 residues in the KRAS complex (B). The binding energy hotspots correspond to residues with high mutational sensitivity. The heatmaps show the computed binding free energy changes for 20 single mutations on the sites of variants. The squares on the heatmap are colored using a 3-colored scale yellow-white-blue-red, with blue color indicating the appreciable unfavorable effects on stability and red pointing to very large destabilizing effects of mutations. The horizontal axis represents the binding epitope residues. We used the BeAtMuSiC contact predictor^60,61^ to identify binding interface residues. Residues are considered part of the interface if they are within a defined cutoff distance 5 Å from atoms in the binding partner. The Y axis depicts all possible substitutions of a given RBD binding epitope residue denoting mutations to letters using a single letter annotation of the amino acid residues.

Monobodies 12D1 and 12D5 demonstrate similar modes of interaction with KRAS(G12D) but reflect their unique structural adaptations to the SWII pocket. While 12D1 induces conformational changes through an enlarged binding footprint, 12D5 achieves high specificity with a more compact interface. The mutational heatmap of the KRAS-12D1 complex revealed several important binding affinity hotspots that correspond to V9, D12, K88, H95, Y96 and I100 residues (Figure 3A). Strikingly, the largest destabilization changes are associated with mutations of H95 and especially Y96 residues (Figure 3A). In particular, we found several highly destabilizing mutations Y96D (ΔΔG = 3.46 kcal/mol), Y96P (ΔΔG = 3.36 kcal/mol), Y96E (ΔΔG = 3.21 kcal/mol) and Y96N (ΔΔG = 3.14 kcal/mol) (Figure 3A). Other notable destabilizing mutations were detected for V9 and H95 residues. Despite the obvious importance of the hydrogen bond formed with the side chain of G12D position on KRAS, mutations in this site produced relatively moderate binding losses (Figure 3A).

On the other hand, mutational profiling highlighted the critical role of H95/Y96 pocket in KRAS that produces the bulk of the binding affinity of the 12D1 MB (Figure 3A). The H95/Y96 pocket is an allosteric site formed by residues H95 and Y96 in the α3 helix of KRAS. These sites are immediately adjacent to the SWII pocket which is the primary binding site for both 12D1 and 12D5. These results of mutational scanning are in excellent agreement with the experimental studies^31^ showing that H95/Y96 allosteric pocket in KRAS is a critical region that has emerged as a hotspot for selective targeting of oncogenic KRAS mutants, including KRAS-G12D. (Figure 3A). Mutational heatmap of the monobody 12D1 residues highlighted the key binding hotspots V30, F31 and triad of adjacent residues L77, W78, and Y79 (Figure 3B). The mutational scanning particularly pointed to the key residues of the FG loop of 12D1 (residues 78–82) with residues W78, Y79, S80, G81, and Y82 occupying space normally occupied by the SWII region in unbound KRAS. According to our results, L77, W78, and Y79 make the largest specific contribution, while the impact of S80, G81 and Y82 appeared to be somewhat less critical as mutations in these positions are less dramatic (Figure 3B). Interestingly, the backbone NH group of G81 on 12D1 makes hydrogen bond with the side chain of D12, while the backbone of S80 forms multiple hydrogen bonds with A59 and Q61 on KRAS, stabilizing the interface and contributing to binding affinity. It is important to note that the knowledge-based energy function employed for rapid mutational profiling of KRAS binding aggregates the effects of protein stability and intermolecular interactions. As a result, the largest effects are seen for monobody residues that are important for both folding stability and binding with KRAS, i.e. L77, W78 and Y79 sites (Figure 3B).

Unlike 12D1, 12D5 occupies a more compact volume within the SWII pocket, resulting in a smaller displacement of the SWII region. 12D5 also interacts with the H95/Y96 pocket but its engagement is likely less extensive compared to 12D1 due to its more compact binding mode. We found that the H95/Y96 pocket remains important for the selectivity of 12D5 (Figure 3C) as it provides a unique recognition site that distinguishes KRAS(G12D) from other RAS isoforms.^31^ Mutational heatmap for 12D5 binding with KRAS is similar to that of 12D1, revealing critical hotspots at positions V9, D12, H95, Y96 and I100 (Figure 3C). In particular, the highly destabilizing mutations were seen for Y96D (ΔΔG = 3.26 kcal/mol), Y96P (ΔΔG = 3.26 kcal/mol), along with V9E (ΔΔG = 3.22 kcal/mol), V9D (ΔΔG = 3.01kcal/mol), V9P (ΔΔG = 2.94 kcal/mol), Y96E (ΔΔG = 2.94 kcal/mol), and Y96N (ΔΔG = 2.93 kcal/mol) (Figure 3C). Interestingly, mutational heatmap for 12D5 binding with KRAS similarly revealed that residues V9 and I100 in KRAS play important roles in the binding interactions although their contributions are secondary compared to H95 and Y96 (Figure 3C). V9 contributes to 12D1/12D5 binding by stabilizing the conformation of the P-loop, which interacts with residues in the monobody’s binding interface. More specifically, V9 of KRAS makes favorable hydrophobic interactions with L77, W78 residues of 12D1 and 12D5 (Figure 3C). V9 contributes to the structural integrity of the P-loop ensuring proper positioning of residues that engage in direct interactions with monobodies. I100 is located near the H95/Y96 allosteric pocket and interacts with L78 and W78 residues of 12D5 monobody (Figure 3C). While V9 and I100 are not primary determinants of monobody binding, they play supporting roles by stabilizing critical regions of KRAS that are directly involved in interactions with monobodies. I100 stabilizes the α3 helix and supports the H95/Y96 allosteric pocket, enhancing the overall binding affinity and selectivity of monobodies. Mutational profiling of 12D1/12D5 binding with KRAS-G12D suggested cooperative stabilization showing that binding to the H95/Y96 pocket can work synergistically with interactions at the SWII pocket and D12 of KRAS to stabilize the overall complex. Mutational heatmap of the monobody 12D5 is similar to the corresponding map of 12D1 residues revealing binding hotspots L77, W78, and Y79 (Figure 3D). We suggest that the H95/Y96 pocket is a key determinant of both binding and selectivity because residues in this pocket differ subtly between KRAS and other RAS isoforms, allowing for isoform-specific recognition.^31^ The critical KRAS binding hotspots form interactions with monobody hotspots V30, F31, and L77. At the same time, the triad L77-W78-Y79 of monobodies form a network of interactions with a wide range of KRAS residues (57-70) imposing significant binding contribution. The interactions of monobodies with G12D position involve residue K75, Y76 S80, G81 and Y82 (Figure 3B,D). Interestingly, these specific contacts targeting D12 sites are formed by another group of monobody hotspots (S80, G81, Y82).

Structural mapping of the binding energy hotspots for the KRAS-G12D complexes with monobodies illustrated the critical role of G12D position and especially residues H95/Y95 in the pocket between SWII region and the α3 helix of KRAS (Figure 4). These sites are immediately adjacent to the SWII pocket which is the primary binding site for both 12D1 and 12D5. The results of mutational scanning are consistent with the experimental studies^31^ which demonstrated the importance of H95 for binding. Indeed, it was found that the introduction of the Q95H mutation in HRAS to mimic KRAS promoted the binding of 12D1 to HRAS. Our results also captured the main recognition features of KRAS-G12D found in these experiments, i.e. G12D mutation, the GTP-bound state, and presence of H95 position as critical binding hotspot.^31^ Notably, binding o KRAS-G12C with s inhibitor sotorasib (AMG-510) similarly exploits H95 as a critical hotspot driving specific recognition and binding affinity.^85^

**Figure 4.**
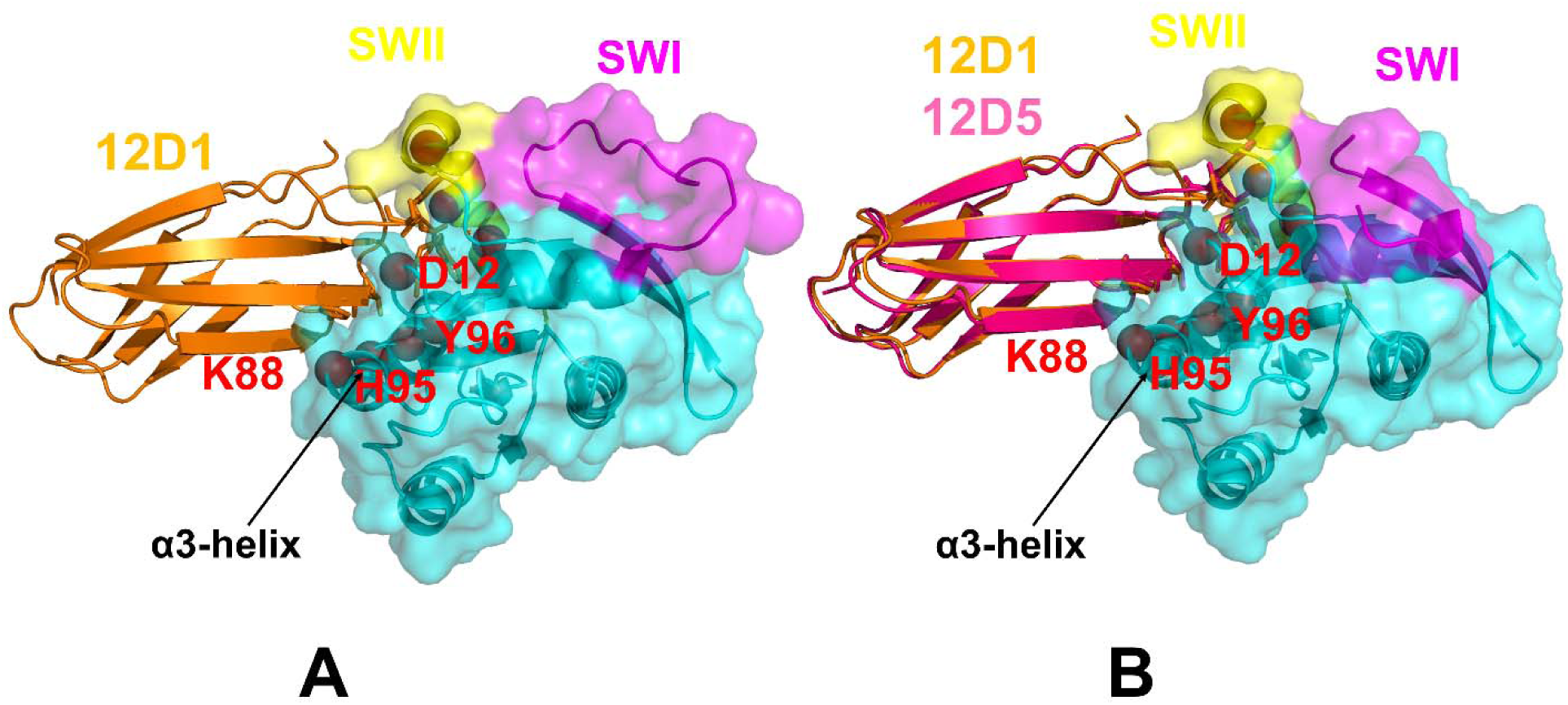
Structural mapping of the binding energy hotspots revealed in mutational scanning for KRAS-G12D protein complexes with monobodies 12D1/12D5. (A) Structural mapping of the binding energy hotspots (show in red spheres) on the crystal structure of KRAS-G12D protein with monobody 12D1 (pdb id 8EZG). The highlighted functional regions of KRAS are annotated: switch I, SW1 (residues 24-40 in magenta ribbons), switch II, SWII (residues 60-76 in yellow ribbons), and the α-helix 3 (residues 87-104 in red ribbons). KRAS protein is shown in cyan ribbons and transparent surface. The monobody 121 is in orange ribbons. The key binding hotspots on KRAS-G12D (D12, H95 and Y96) are annotated. (B)) Structural overlay of the crystal structures of KRAS-G12D with monobody 12D1 (in orange ribbons) and 12D5 (in hotpink ribbons). The highlighted functional regions of KRAS (SWI, SWII and α-helix 3) are annotated as in panel (A).

### Probing KRAS Binding Interactions with Affimer Proteins Through Mutational Scanning Reveals Role of Reciprocal Binding Energy Hotspots for KRAS and Binding Partners

We also performed mutational profiling of KRAS binding with affimer proteins K6, K3 and K69 (Figure 5). The mutational heatmap of the KRAS-K6 complex revealed several important binding affinity hotspots D33, I36, L56, M67, Y71 and T74 (Figure 5A). The importance of these amino acid residues for K6 function was confirmed by mutational analysis.^38^ Our mutational scanning revealed the key role of L56 and Y71 positions (Figure 5A). The largest destabilization changes are associated with mutations I36P (ΔΔG = 2.69 kcal/mol), M67D (ΔΔG = 2.41 kcal/mol) and Y71P (ΔΔG = 2.12 kcal/mol). I36 interacts with affimer K6 hydrophobic positions F40, M49 and W70, while L56 binds with W43, P42 and M67 binds with I74, K72 and W43 (Figure 5A).

**Figure 5.**
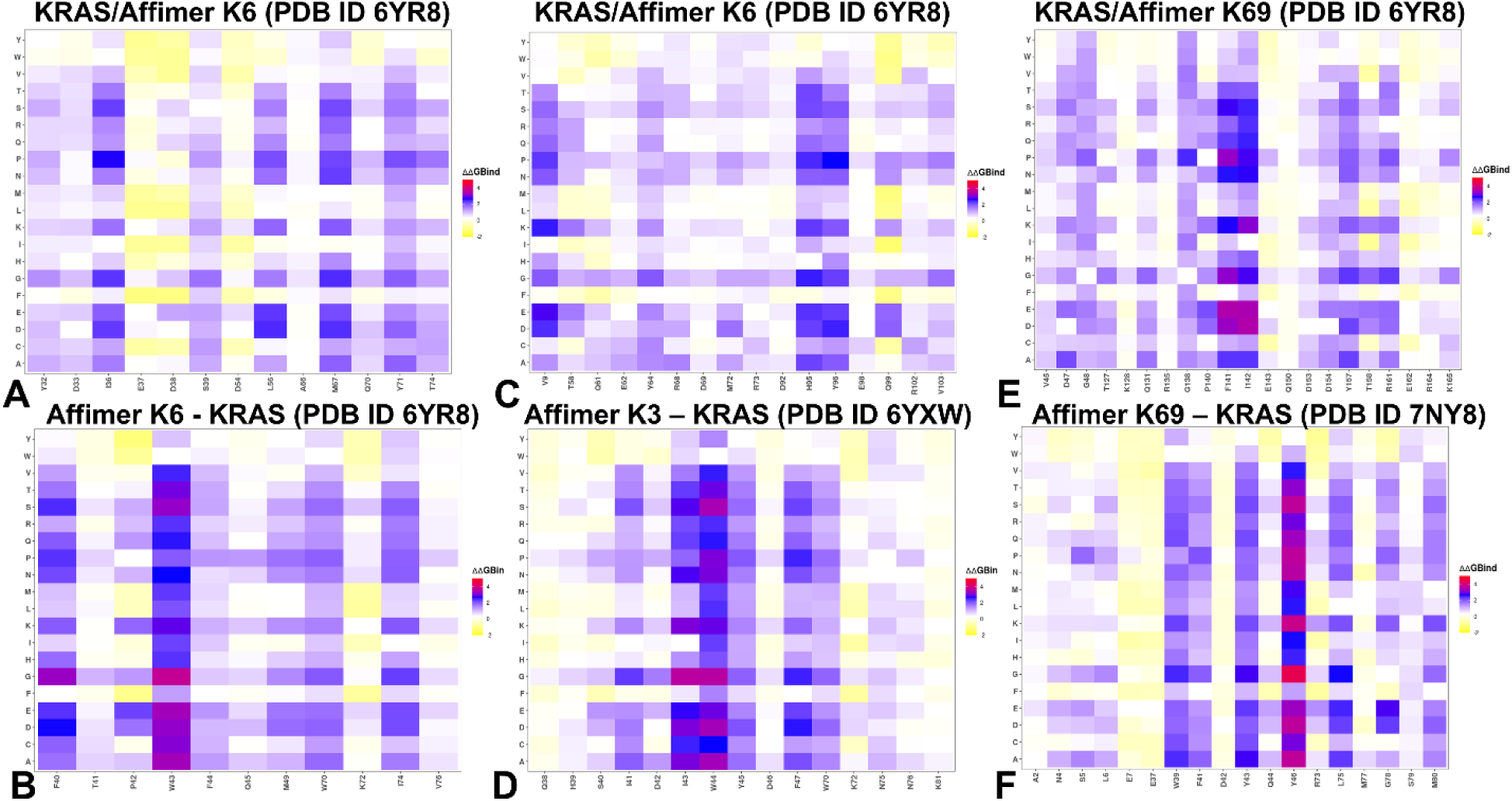
The ensemble-based mutational scanning of binding for the KRAS complexes with affimer proteins K6, K3 and K69. The mutational scanning heatmap for the KRAS binding epitope residues in the KRAS complex with affimer K6 (A) and mutational heatmap for affimer K6 (B). The mutational scanning heatmap for the KRAS binding epitope residues in the KRAS complex with affimer K3 (C) and mutational heatmap for affimer K3 (D). The mutational scanning heatmap for the KRAS binding epitope residues in the KRAS complex with affimer K69 (E) and mutational heatmap for affimer K69 (F). The binding energy hotspots correspond to residues with high mutational sensitivity. The heatmaps show the computed binding free energy changes for 20 single mutations on the sites of variants. The squares on the heatmap are colored using a 3-colored scale yellow-white-blue-red, with blue color indicating the appreciable unfavorable effects on stability and red pointing to very large destabilizing effects of mutations. The horizontal axis represents the RBD binding epitope residues. Residues are considered part of the interface if they are within a defined cutoff distance 5 Å from atoms in the binding partner. The Y axis depicts all possible substitutions of a given RBD binding epitope residue denoting mutations to letters using a single letter annotation of the amino acid residues.

Mutational scanning of affimer K6 interface residues highlighted the importance of residues 40–45 from variable region 1 (Figure 5B). The affimer tripeptide motif formed of P42, W43, and F44 binds the shallow hydrophobic pocket of KRAS. We found that W43 is the central binding hotspot on K6 since all mutations of W43 lead to large destabilizing changes (Figure 5B). W43 forms a large number of favorable contacts with many KRAS positions including K5, L6, V7, S39, D54, I55, L56, M67, Q70, Y71 and T74. Importantly, mutational scanning maps showed that the hydrophobic cluster formed by W43 of affimer K6 and V7/I55/L56/M67 sites of KRAS is central for binding (Figure 5B). Our results are consistent with the importance of these amino acid residues revealed by mutational analysis and functional studies showing that replacements of P42, W43, F44, or Q45 with alanine reduced affimer-mediated inhibition of nucleotide exchange.^38^ Mutational scanning of KRAS binding interface residues showed the larger binding epitope for K3, revealing key hotspot positions V9, Y64, M72 and especially H95 and Y96 (Figure 5C).

The respective mutational scanning of affimer K3 interfacial residues showed critical hotspots at positions 40-45 in variable region I (VRI) particularly I43, W44, Y45 and F47 (Figure 5D). H95 of KRAS makes multiple contacts with Y45, N76, L74, N75, I43, W44 of affimer K3 while Y96 is engaged in favorable interactions with I43 and W44 of K3. The importance of these interactions was correlated with pulldown and nucleotide exchange assays in which residues I41, D42, I43, W44, Y45, and D46 when mutated to alanine, did not bind and inhibit KRAS.^38^ The mutual correspondence between KRAS hotspots H95/Y96 and K3 hotspots I43/W44 provides convincing evidence that the interaction network formed by these residues is central to the binding specificity (Figure 5C,D).

These results are supported by biochemical and cellular data showing that binding W44 with KRAS H95 may explain the specificity of K3 for KRAS as H95 is a unique residue only present in KRAS and not in HRAS and NRAS^38^. The importance of H95 in mediating affimer K3 selectivity for KRAS was confirmed by immunoprecipitation assays of mutants H95Q and H95L that mimic the corresponding residues in HRAS and NRAS.^38^ According to our data, mutations H95Q (ΔΔG = 2.83 kcal/mol) and H95L (ΔΔG = 3.25 kcal/mol) produced large destabilization changes and loss of binding (Figure 5C). Hence, the results of mutational scanning based on conformational ensembles produced excellent agreement with the experimental data, accurately reproducing the functional hotspots of both KRAS and affimer proteins.

Unlike affimers K3 and K6, K69 offers a complementary mechanism for inhibiting KRAS signaling through an indirect, long-range mechanism. Mutational scanning revealed a range of hotspots, most prominently featuring P140, F141, I142, and Y157 sites where mutations would disrupt the hydrophobic core, destabilizing the complex (Figure 5E). Mutational scanning showed that Y157 and R161 are also important binding hotspots, albeit producing less dramatic changes upon mutations as compared to the triad P140/F141/I142. At the same time, mutational scanning of affimer K69 binding residues identified notable hotspots at positions W39, F41, Y43, Y46 and L75 (Figure 5F). Both Y43 and Y46 of affimer K69 participate in interactions with P140 and F141 of KRAS forming a tight hydrophobic cluster that determines the binding and selectivity profile of affimer K69 inhibition.

Structural mapping of the binding energy hotspots determined by mutational scanning highlighted the distribution of the binding epitopes and location of major energetic centers for affimer proteins (Figure 6). The major hotspots for K6 include I36, L56, M67 (Figure 6A), for K3 (Y64, H95, Y96 and Q99) (Figure 6B). Affimer K6 binds to a pocket on KRAS between the SWI and SWII regions, while affimer K3 binds to the SWII/α3 helix pocket. Affimer K69 binds the allosteric lobe between helices 4 and 5 on the opposite side of KRAS to affimers K3 and K6 (Figure 6C). The location of the binding pockets and distribution of the binding energy hotspots highlighted the role of SWI, SWII, and α3 helix as critical functional regions for KRAS binding. Importantly, the structural mapping of the KRAS binding hotspots in the allosteric lobe (Figure 6C) showed that K69 binding may induce allosteric communication to the other functional regions of KRAS, thus potentially linking distinct binding sites in a cooperative interaction network that can be modulated by different affimer proteins.

**Figure 6.**
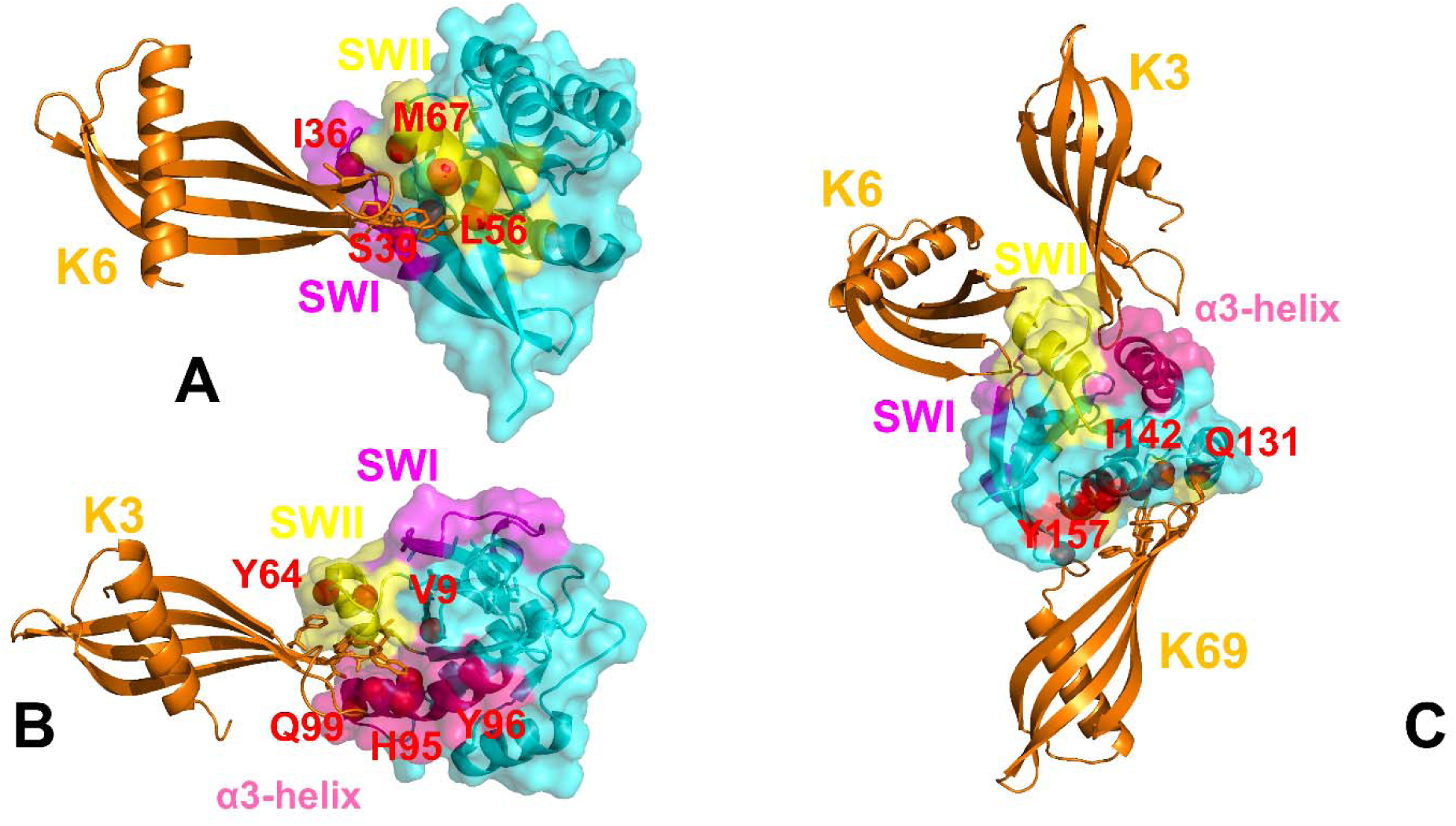
Structural mapping of the binding energy hotspots in the KRAS complexes with affimer proteins K6, K3 and K69. (A) Structural projection of binding energy hotspots (red spheres) on the crystal structure of KRAS protein with K6 (pdb id 6YR8). KRAS proteins are shown in cyan ribbons and transparent surface. Affimer K6 is in orange ribbons and annotated. The highlighted functional regions of KRAS are annotated: SW1 (residues 24-40 in magenta ribbons), SWII (residues 60-76 in yellow ribbons). The KRAS hotspots I36, S39 L56, M67 are in spheres and annotated. (B) Structural projection of binding energy hotspots (red spheres) on the crystal structure of KRAS protein with K3 (pdb id 6YXW). Affimer K3 is shown in orange ribbons. The annotation of SWI and SWII regions is the same as on panel (A). Additionally, α-helix 3 is shown in red ribbons. The binding hotspots V9, Y64, H95, Y96 and Q99 are in red spheres and annotated. (C) Structural projection of binding energy hotspots (red spheres) on the crystal structure of KRAS with K69 (pdb id 7NY8). The crystal structures of KRAS with affimer proteins are overlayed and all three affimer proteins are shown in orange ribbons. The binding hotspots of KRAS for binding with K69 (residues Q131, I142, I157) are in red spheres and annotated.

To summarize, despite KRAS being historically considered “undruggable,” monobodies and affimer proteins induce formation of druggable pockets. The SWII pocket (SII-P) and H95/Y96 pocket are examples of cryptic allosteric sites that become accessible during conformational fluctuations. Monobodies and affimers exploit these transiently exposed pockets, stabilizing specific conformations and modulating KRAS activity. Mutational profiling of KRAS complexes with affimer proteins highlighted the diversity of binding mechanisms and a wide range of potential hotspots particularly pointing to the unique role of the SII/α3 pocket by targeting H95/Y96 sites stabilizing a cryptic conformer and inhibiting nucleotide exchange. The results of dynamic and energetic analysis underscores that each affimer employs a distinct mechanism to stabilize inactive states. Affimer K6 stabilizes the SI/SII hydrophobic pocket, preventing transitions to active states. Affimer K3 targets the cryptic SII/α3 pocket, leveraging the α3 helix as a hinge point to amplify its effects while affimer K69 induces global conformational changes by targeting the allosteric lobe, acting as a hub for long-range communication. The analysis of KRAS binding with monobodies and affimer proteins suggested that targeting hotspots can induce allosteric perturbations that propagate through the protein structure, affecting distant regions. In particular, targeting the α3-helix and allosteric lob regions induces global changes, destabilizing effector interactions and nucleotide exchange. These allosteric effects can amplify the functional impact of targeting hotspots, making them attractive therapeutic targets.

### Binding Free Energy Analysis of the KRAS Complexes with Monobodies and Affimer Proteins Quantifies the Energetic Drivers of Binding and Specificity

Using the conformational equilibrium ensembles obtained MD simulations of the KRAS complexes we computed the binding free energies for these complexes using the MM-GBSA method.^72–75^ MM-GBSA is also employed to (a) decompose the free energy contributions to the binding free energy of a protein–protein complex on per-residue basis; (b) evaluate the role of hydrophobic and electrostatic interactions as thermodynamic drivers of KRAS binding; (c) identify the binding energy hotspots and compare the rigorous MM-GBSA predictions of binding energetics with the results of mutational scanning in the KRAS complexes.

We analyzed the MM-GBSA results for the KRAS complex with monobodies 12D1 (Figure 7A-C) and 12D5 (Figure 7D-F). The energy decomposition showed that the binding affinity hotspots for 12D1 correspond to KRAS residues K88 (ΔG = −6.17 kcal/mol) followed by mutations Q61 (ΔG = −5.21 kcal/mol), Y64 (ΔG = −4.44 kcal/mol), Y96 (ΔG = −3.97 kcal/mol) and H95 (ΔG = −3.79kcal/mol) (Figure 7A). The contribution of van der Waals interactions is most favorable for H95 (ΔG_VDW_ = −6.41 kcal/mol), Y64 (ΔG_VDW_ = −5.13 kcal/mol) and Y96 (ΔG_VDW_ = −4.59 kcal/mol) (Figure 7B), while the electrostatic contribution drives binding of K88 (ΔG_ELE_ = −81.39 kcal/mol), D69 (ΔG_ELE_ = −21.84 kcal/mol) and G12D mutated position (ΔG_ELE_ = −14.71 kcal/mol) (Figure 7C). These results reflected the presence of stable hydrogen bonding formed by K88 of KRAS with D33 and E47 of 12D1. K88 also makes contacts with important hotspots on 12D1 such as F31, T49 and Y82. Q61 on KRAS makes specific interactions with monobody hotspots Y79, S80 and Y82 which contributes to the highly favorable total binding energy (Figure 7A-C). The results of mutational scanning showed that residues W78, Y79, S80, and Y82 are the most important hotspots of 12D1. As a result, MM-GBSA analysis suggests that the specific interactions of K88 and Q61 formed with this hotspot cluster make decisive contributions to the binding. MM-GBSA analysis confirmed the importance of H95/Y96 positions for both stability and binding. 12D1 engages the H95/Y96 pocket through specific interactions formed by the side chain of F31 in 12D1 making π–π stacking interactions with H95 and the side chain of T49 interacting with H95 via a hydrogen bond, further enhancing binding affinity. These interactions contribute to the stabilization of the enlarged SWII pocket conformation induced by 12D1 monobody.

**Figure 7.**
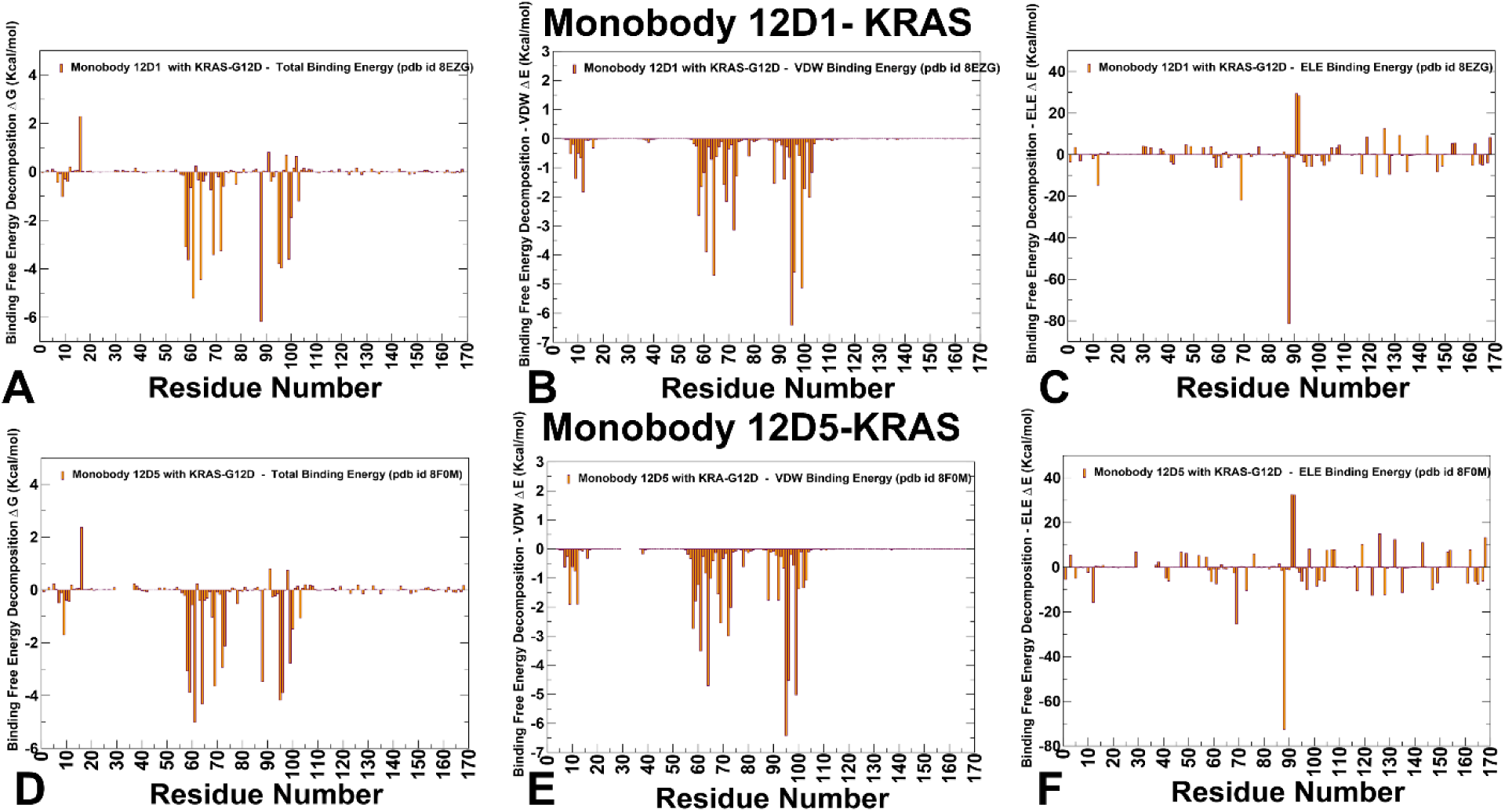
The residue-based decomposition of the total binding MM-GBSA energies for the KRAS residues in the KRAS-G12D complex with monobody 12D1 (A-C) and monobody 12D5 (D-F). The residue-based decomposition of the total binding energy (A), van der Waals contribution (B) and electrostatic contribution to the total MM-GBSA binding energy (C) for the KRAS residues in the KRAS-G12D complex with 12D1. The residue-based decomposition of the total binding energy (D), van der Waals contribution (E) and electrostatic contribution to the total MM-GBSA binding energy (F) for the KRAS residues in the KRAS-G12D complex with 12D5. The MM-GBSA contributions are evaluated using 1,000 samples from the equilibrium MD simulations of respective KRAS complexes. It is assumed that the entropy contributions for binding are similar and are not considered in the analysis.

A generally similar breakdown of the energetic hotspots and contributions emerged from MM-GBSA analysis of KRAS-12D5 complex where the dominant total contributions were seen for Q61 (ΔG = −5.01 kcal/mol), Y64 (ΔG = −4.32 kcal/mol), H95 (ΔG = −4.17 kcal/mol) and Y96 (ΔG = −3.89 kcal/mol) (Figure 7D). The results confirmed that the van der Waals contribution drives binding of H95 and Y96 hotspots with (ΔG_VDW_ = −6.42 kcal/mol) for H95 and (ΔG_VDW_

= −4.52 kcal/mol) for Y96 hotspots (Figure 7E). Similar to 12D1, the electrostatic contributions are the largest for K88 (ΔG_ELE_ = −72.63 kcal/mol), D69 (ΔG_ELE_ = −25.39 kcal/mol) and D12 (ΔG_ELE_ = −15.7 kcal/mol) (Figure 7F). Combined, the results of mutational scanning and MM-GBSA computations established that KRAS binding with monobodies 12D1/12D5 is largely driven by a handful of major binding hotspots such as Q61, K88, H95 and Y96 that are for the most part are located in the α3 helix (residues 87-104) that is selectively targeted by monobodies.

MM-GBSA analysis of the KRAS binding with affimer proteins enabled a detailed comparison with mutational scanning data and more rigorous assessment of the energetic contributions and primary binding hotspots (Figure 8). We found that binding to affimer K6 is driven primarily by SWI hotspots I36, D38, S39 as well as SWII residues L56, M67, Q70 and Y71 (Figure 8A). The largest total contributions were seen for I36 (ΔG = −4.27 kcal/mol), S39 (ΔG = −3.47 kcal/mol), L56 (ΔG = −3.47 kcal/mol) and Y71 (ΔG = −2.32 kcal/mol) (Figure 8A). These results are consistent with the mutational scanning analysis showing the central role of L56 and Y71 sites as well as in line with the experimental studies.^38^ Indeed, the predicted KRAS binding hotspots are targeted by a hydrophobic cluster of K6 that bind to I36, L56, M67, and Y71 sites on KRAS from SWI and SWII regions. Consistent with structural studies^38^, the binding hotspots are mainly determined by highly favorable hydrophobic contacts and van der Waals interactions (Figure 8A,B). The favorable electrostatic interactions are seen for SWI residues D33, E37, D38 as well as D54 and D69 positions (Figure 8C). Notably, despite appreciable contributions from these residues, none of these KRAS positions are among major binding hotspots. These findings highlighted the central role of hydrophobic packing in the driving binding of affimer K6.

**Figure 8.**
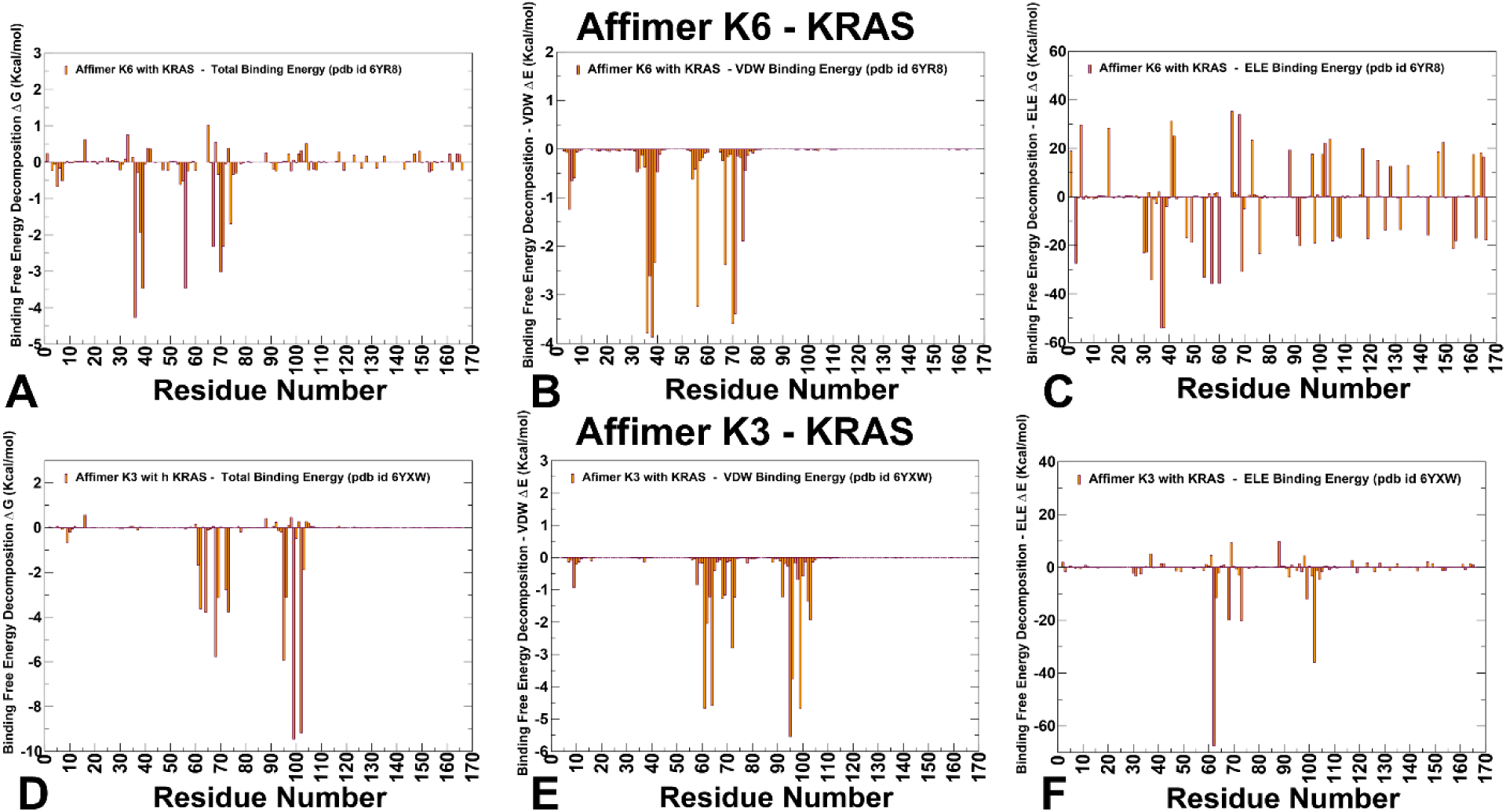
The residue-based decomposition of the total binding MM-GBSA energies for the KRAS residues in the KRAS complex with affimer K6 (A-C) and affimer K3 (D-F). The residue-based decomposition of the total binding energy (A), van der Waals contribution (B) and electrostatic contribution to the total MM-GBSA binding energy (C) for the KRAS residues in the KRAS complex with affimer K6. The residue-based decomposition of the total binding energy (D), van der Waals contribution (E) and electrostatic contribution to the total MM-GBSA binding energy (F) for the KRAS residues in the KRAS complex with affimer K3. The MM-GBSA contributions are evaluated using 1,000 samples from the equilibrium MD simulations of respective KRAS complexes.

Affimer K3 binds to the cryptic SWII/α3 pocket, a region that includes the SII region (residues 60-76) and the α3-helix (residues 87–104).^38^ MM-GBSA computations provided a more granular characterization of binding hotspots with K3 showing the critical role of Q99 (ΔG = −9.45 kcal/mol) and R102 (ΔG = −9.17 kcal/mol) along with previously detected hotspots H95 (ΔG = −5.92 kcal/mol) and Y96 (ΔG = −3.12 kcal/mol) (Figure 8D). MM-GBSA analysis showed that most favorable hydrophobic contacts are made by H95, Q99, Q61, Y64, Y97, M72, and V103 (Figure 8E). At the same time, the electrostatics dominates contributions of R102, R73, R68 and Q99 (Figure 8F). When affimer K3 binds to the SII region, the D42 residue of K3 brings the SII region close to the α-3 helix by generating a hydrogen bond between R68 of SII and Q99 of α-3 helix. The complex formation is further strengthened by hydrogen bonds between side chain oxygen of D46 residue in K3 with the main chain nitrogens of Q99 and R102 in the α3-helix region of KRAS. This hydrogen bonding network anchors binding interfaces where I41, I43, W44 of K3 form hydrophobic interactions with V9, M72, V103, and H95 of KRAS (Supporting Information, Figure S3). Overall, MM-GBSA analysis provided important insights into nature of binding mechanism of KRAS with affimer K3 that is anchored by specific interactions of Q99 and R102 hotspots that enable hydrophobic packing KRAS hotspots H95/Y96 and K3 hotspots I41/I43/W44.

MM-GBSA analysis of KRAS binding with affimer K69 revealed major contributions to binding from Q131, I142, F141, P140, Y157 and R161 (Supporting Information, Figure S4A). These results are consistent with the mutational scanning using simplified energy model that similarly precited P140, F141, I142, Y157 and R161 sites. The hydrophobic interactions favor F141, P140, Q131 sites (Supporting Information, Figure S4B), while electrostatics favored contributions of D154, D153, D47, E143 and D132 sites (Supporting Information, Figure S4C).

To conclude, MM-GBSA analysis of KRAS binding with monobodies (12D1, 12D5) and affimer proteins (K6, K3, K69) revealed several key findings regarding the energetic contributions, binding hotspots, and mechanisms driving these interactions:. For both 12D1 and 12D5, the dominant binding hotspots are located in the α3 helix (residues 87–104), specifically Q61, K88, H95, and Y96. While the van der Waals interactions drive binding for H95 and Y96, electrostatic contributions dominate for K88 and D69. We also found that hydrophobic interactions play an important role in binding, with affimer K6 targeting a hydrophobic cluster in KRAS’s SWI/SWII regions, while K3 binding is stabilized by a hydrogen-bonding network and hydrophobic packing between KRAS hotspots (H95/Y96) and K3 hotspots (I41/I43/W44). Affimer proteins target distinct regions of KRAS (SWI/SWII for K6, cryptic SWII/α3 pocket for K3, allosteric lobe for K69), highlighting their diverse binding mechanisms.

Overall, MM-GBSA studies suggested that KRAS binding with monobodies and affimer proteins is driven by specific hotspot residues (e.g., Q61, K88, H95, Y96) and are often dominated by hydrophobic interactions, with additional important contributions from electrostatics and hydrogen bonding that provide recognition anchors of specificity. MM-GBSA results align well with mutational scanning and experimental data, validating the method’s ability to predict binding energetics and identify hotspots.

### Mutational Profiling of Residue Interaction Networks Maps Allosteric landscape and Hotspot Clusters of Allosteric Communications

We used network centrality analysis and the network-based mutational profiling of allosteric residue propensities that are computed using topological network parameter ASPL to characterize global network of allosteric communications. Through ensemble-based averaging over mutation-induced changes in these network metric, the proposed model can identify positions in which mutations on average cause global network changes and correspondingly alter allosteric interactions and long-range communications in KRAS. Allosteric hotspots are identified as residues in which mutations incur significant edgetic perturbations of the global residue interaction network that disrupt the network connectivity and cause a significant impairment of global network communications and compromise signaling. This analysis enables identification of allosteric control points that could determine the efficient and robust long-range communications in the KRAS complexes. In this model, residue nodes with high Z-score ASPL values can function as bottlenecks in the global interaction network, affecting the robustness and efficiency of communication between different parts of the network. We first examined Z-score ASPL profiles for KRAS residues in complexes with monobodies 12D1/12D5 (Figure 9A,B). In the KRAS complex with 12D1, we detected a relatively small number of profound peaks that are distributed across different functional regions of KRAS and correspond to residues 19-23 located near P-loop, D30 from SW1 region, residues R68, M72 (SWII), F78, L80 (β-strand 4), F90, H95, Y96 (α-helix 3) along with positions V114, N116 (β-strand 5) and L133, V152, F156 (from α-helix 4: 127–136 and α-helix 5: 148–166) (Figure 9A). These residues appeared to form contiguous network that links the key binding interface positions (D12, K88, Q61, D69, H95,Y96) to the allosteric lobe through critical for allostery α-helix 3, β-strand 4 and α-helix 5 segments of KRAS (Figure 9A, 10A). As may be expected, the Z-score ASPL profile for KRAS binding with 12D5 revealed similar distribution of peaks and KRAS positions where mutations can alter dramatically the allosteric interactions (Figure 9B). Among prominent profile peaks are residues F78, L79, L80, and F82 from β-strand 4 as well as I93, Y96 from α-helix 3 and V114 and N116 from β-strand 5 that mediate long-range communication from KRAS-12D5 binding interface to Y157, L159, I163 and R164 from α-helix 5 of the allosteric lobe (Figure 9B,10B).

**Figure 9.**
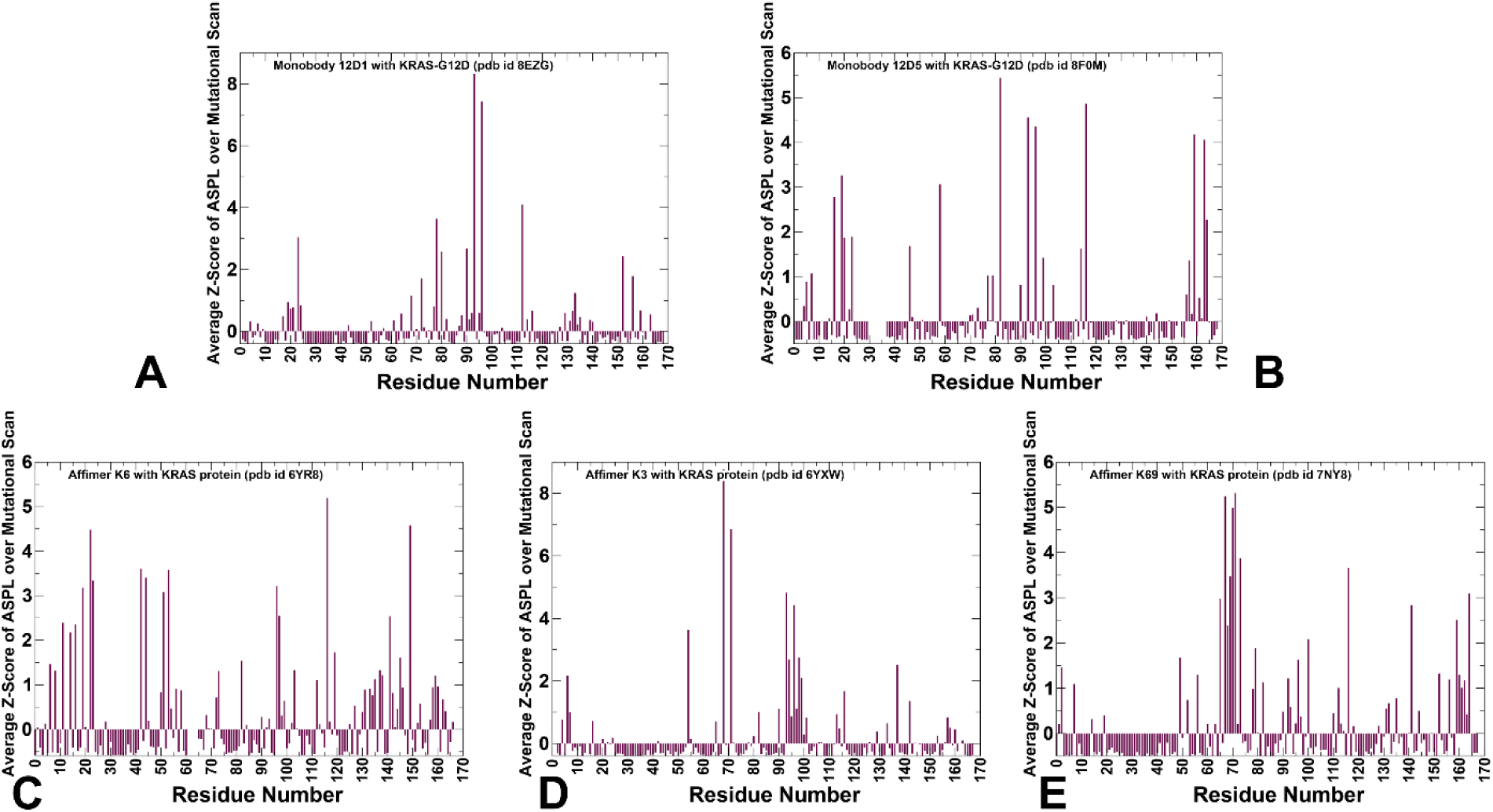
Network centrality analysis of the KRAS complexes with monobodies and affimer proteins. (A) The average Z-score of the ASPL network metric over mutational scan. Z-score is obtained for each node based on the change of the characteristic path length under node removal averaged over all protein residues. The Z-score are then computed over all mutational changes in given position. The network metric profiles are shown for KRAS-G12D complex with monobodies 12D1 (A), 12D5 (B) and KRAS protein complexes with affimer protein K6 (C), affimer K3 (D) and affimer K69 (E). The profiles are shown in maroon-colored filled bars.

**Figure 10.**
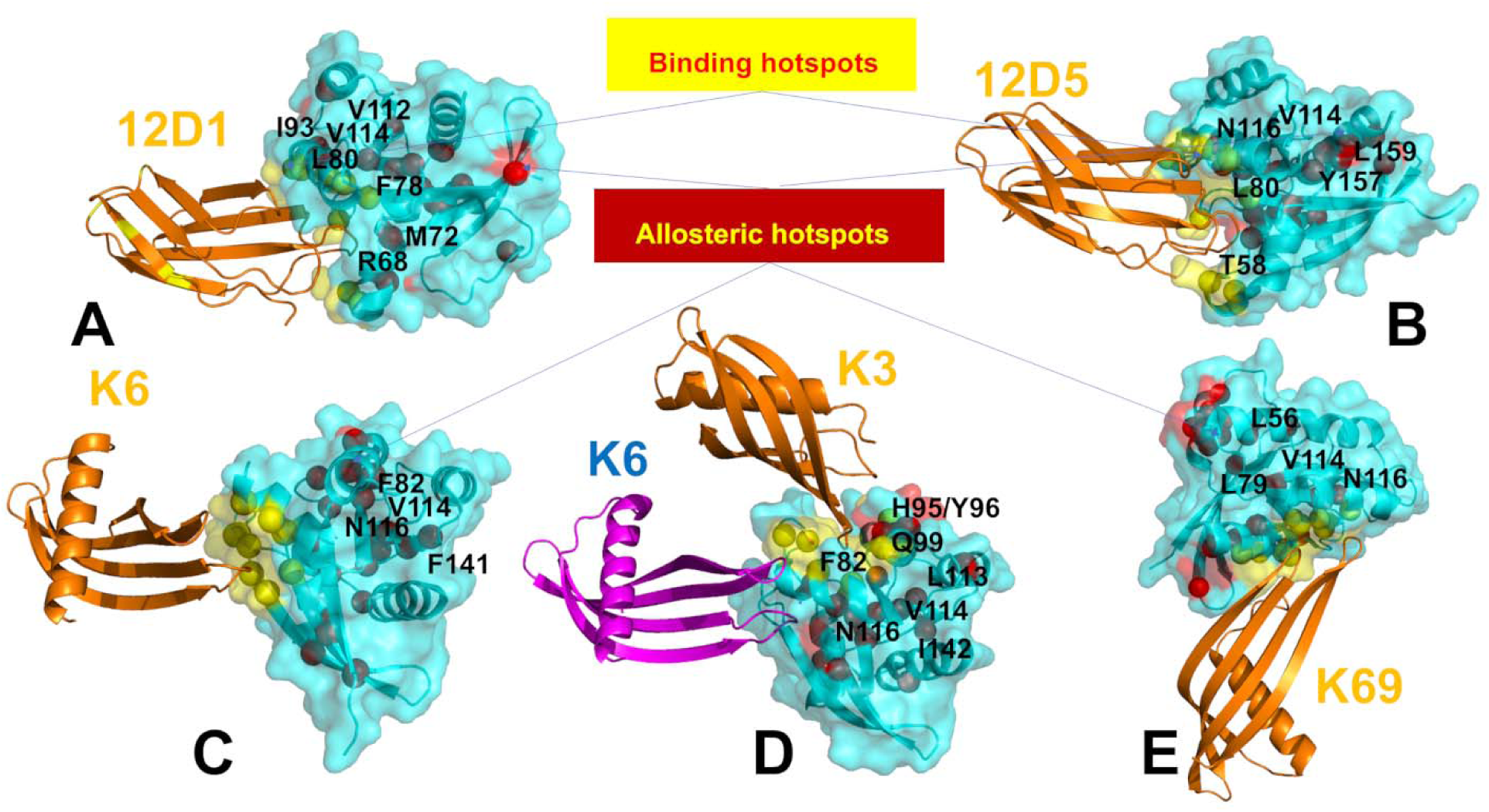
Structural mapping of the allosteric hotspots in the KRAS complexes with monobodies 12D1/12D5 (A,B) and affimer proteins K6, K3 and K69 (C-E). Structural projection of allosteric hotspots (red spheres) and binding energy hotspots (yellow spheres) on the crystal structure of KRAS protein with monobody 12D1 (A) and 12D5 (B). Monobodies 12D1/12D5 are shown in orange ribbons. The allosteric hotspots are annotated. Structural projection of allosteric hotspots (red spheres) and binding energy hotspots (yellow spheres) on the crystal structure of KRAS protein with affimer K6 (C), K3 (D) and K69 (E). KRAS proteins are shown in cyan ribbons and transparent surface. Affimer K6 is in orange ribbons and annotated. On panel (D) the affimer K6 (blue ribbons) is overlayed with the structure of KRAS bound to K3. The predicted allosteric hotspots are in red spheres and major allosteric centers are annotated.

The most important finding of this analysis is that the predicted allosteric hotspots are consistent with and in excellent agreement the seminal experimental studies of KRAS allostery showing that the efficient propagation of allosteric effects is mediated by the central β-strand of KRAS which acts as a hub for transmitting conformational changes linking distant functional sites.^41^ More specifically, the comprehensive DMS analysis of KRAS allostery revealed the important role of KRAS positions F78, G79, F82 from β-strand 4 and residues V112, V114 and N116 from β-strand 5 as allosteric centers affecting binding with effector proteins through long-range interactions. The important revelation of our predictions is the fact that the allosteric binding hotspots may correspond to a conserved group of mediating positions that could control allosteric binding with diverse range of binding partners targeting different pockets on KRAS. In addition, a group of residues from the α-helix 3 (I93, H95 and Y96) is critical for linking the binding interface region with the central β-strand and subsequently to the allosteric lobe. Our results emphasized the role of allosteric pocket located between SWII and α-helix 3 which harbor allosterically important KRAS sites in agreement with the experimental studies.^41^ Finally, the predicted allosterically important positions (Y157, L159, I163 and R164 from α-helix 5) were found in the experimentally validated allosteric pocket located in the C-terminal lobe of the protein (this pocket is formed by residues 97, 101, 107–111, 136–140 and 161–166)and is the most distant pocket from the binding interface.^41^ Our analysis identifies these sites as major hotspots of allosteric communication that affect interaction paths from remote parts of KRAS to the binding interface.

We also performed allosteric network analysis of KRAS binding with affimer proteins (Figure 9C-E). For the KRAS complex with affimer K6, the Z-score ASPL showed peaks at KRAS positions 11-23 located near P-loop, residues K42, V44 (β-strand 2), C51, L53 (β-strand 3), F82 (β-strand 4), Y96 (α-helix 3), V114, N116 (β-strand 5), F141, F143, C145, R149 (β-strand 6) (Figure 8D,9D). These findings supported the notion that the central β-strand acts as an allosteric hub linking distant functional sites.^41^ A different binding interface for KRAS binding with affimer K3 only moderately redistributed the distribution of Z-score ASPL values revealing peaks at important allosteric sites, particularly featuring residues F90, I93, H94, H95, Y96, E98, Q99 (α-helix 3), R68, Y71 (α-helix 2), F82 (β-strand 4) L113, V114 and N116 (β-strand 5), I137, Y142 (β-strand 6) (Figure 9E,10E). Affimer K3 binds to the cryptic SWII/α3 pocket, and we observed that key binding energy hotspots H95, Y96, Q99 can also serve as allosteric centers in the complex (Figure 9E). Interestingly, the network analysis consistently predicted H95/Y96 from SWII/α3 allosteric pocket and V114, N116 (β-strand 5) as allosteric hotspots for both monobodies and affimer proteins. Our results showed that SWII/α3 pocket - which is the binding site of sotorasib and other clinically approved allosteric inhibitors of KRAS - emerged as a conserved allosteric hotspots of long-range communications between KRAS and different binding partners. Interestingly, H95/Y96 residues in this region can be universally important as both binding and allosteric hotspots. The predicted allosteric cluster consists of KRAS residues (H95, Y96, R97) forming cryptic allosteric pocket that is targeted by known allosteric inhibitor sotorasib (AMG 510) which is the first KRAS(G12C) inhibitor to reach clinical testing in humans.^16^ Indeed, structural studies showed that AMG-510 occupied the H95 groove engaged in a network of primarily van der Waals contacts extending from the backbone of helix 2 (H95, Y96) to the flexible switch II loop (Supporting Information, Figure S5). The recent illuminating study from Shokat lab used structural and pharmacological analysis of members of the Ras, Rho, and Rab family of GTPases to demonstrate that this cryptic allosteric pocket in the switch II region can be present in other GTPases indicating that GTPases exhibit targetable allosteric binding pockets.^86^

Finally, the network analysis of KRAS binding with affimer K69 that binds to the allosteric lobe of KRAS away from the binding epitope of K6 and K3 showed a fairly similar distribution of Z-score ASPL peaks featuring residues M67, Q70, Y71, M72, R73 (α-helix 2), Y96 (α-helix 23), L79, F82 (β-strand 4) L113, V114 and N116 (β-strand 5), F141, Y142 (β-strand 6), V152, L159, V160, R164 (α-helix 5) (Figure 9F,10F).

The important finding of this analysis is that despite completely different binding interface with affirmer K69, the predicted group of allosteric hotspots remains relatively conserved, particularly pointing to KRAS positions Y96 (α-helix 3), F82 (β-strand 4), V114 and N116 (β-strand 5), F141 and Y142 (β-strand 6) (Figure 9D-F). These allosteric centers emerged as distribution peaks of the network profile for all KRAS complexes with affimer proteins. Importantly, the significance of these KRAS positions as allosteric binding hotspots was experimentally observed in DMS studies of KRAS binding with six different binding partners that did not include the monobodies and affimer proteins examined here.^41^

To conclude, the analysis reveals a set of highly conserved residues that serve as critical nodes in the allosteric communication network of KRAS. Notably, residues in β-strand 4 (F78, L80, F82), α-helix 3 (I93, H95, Y96), β-strand 5 (V114, N116), and α-helix 5 (Y157, L159, R164) consistently emerge as hotspots across all KRAS complexes studied. These residues form contiguous networks linking the binding interface to the distal C-terminal allosteric lobe, with β-strand 4 acting as a central hub for propagating conformational changes—a finding that strongly corroborates experimental evidence highlighting the role of β-strand 4 as a key mediator of allostery. The SWII/α3 pocket, particularly residues H95 and Y96, is identified as a universal hotspot for both binding and allosteric regulation. This pocket, targeted by clinically approved inhibitors like sotorasib, emerges as a conserved control point for long-range communication between KRAS and diverse binding partners. The predicted hotspots are in excellent agreement with experiments,^41^ which independently identified residues in β-strand 4 (F78, G79, F82) and β-strand 5 (V112, V114, N116) as critical mediators of long-range interactions affecting effector protein binding. Additionally, the C-terminal allosteric pocket (residues 161–166), experimentally validated as a key site for disrupting KRAS signaling, was accurately predicted as a major hotspot in this analysis. This strong alignment between computational predictions and experimental data highlights the robustness of the network-based approach. Despite distinct binding interfaces, shared hotspots (e.g., Y96, F82, V114, N116) highlight a conserved allosteric infrastructure. For instance, affimer K69, which binds far from the epitopes of K6 and K3, still converges on the same critical residues, reinforcing their universal importance in KRAS signaling. By pinpointing residues that function as bottlenecks in the global interaction network, this study not only enhances our understanding of KRAS allostery but also demonstrates the utility of network-based approaches in predicting druggable sites.

## Discussion

The exploration of KRAS dynamics and its interactions with monobodies (12D1, 12D5) and affimer proteins (K6, K3, K69) provides insights into the mechanisms governing KRAS inhibition. The differential effects of monobodies and affimer proteins on KRAS dynamics reveal intriguing insights into the interplay between structural rigidity and functional flexibility. For example, 1rigidifies the SWII region, leading to compensatory flexibility in SWI. In contrast, 12D5 stabilizes both SWII and the α3 helix, suppressing functional movements of SWI. These observations suggest that the balance between rigidity and flexibility is finely tuned, with each binding partner inducing a unique dynamic signature. An interesting observation of our analysis of KRAS dynamics and binding is the interplay between allosteric coupling and functional dynamics in KRAS specifically modulated by binding partners. The stabilization of specific regions such as the SWII pocket and α3 helix by these binding partners induces compensatory flexibility in other regions, particularly the SWI region. For instance, monobody 12D1 stabilizes the SWII region, disrupting the normal coupling between SWI and SWII and leading to enhanced mobility in SWI. This compensatory mechanism likely reflects an attempt by KRAS to maintain functional adaptability despite the immobilization of key regions.

Our results underscored the role of cryptic pockets, such as the H95/Y96 and SII/α3 pockets, in modulating KRAS activity. These transiently exposed sites become accessible during conformational fluctuations and serve as druggable hotspots that stabilize inactive conformations. Monobodies and affimers exploit these pockets to induce specific conformations that inhibit KRAS signaling. For example, affimer K3 targets the cryptic SII/α3 pocket, leveraging the α3 helix as a hinge point to amplify its effects. Similarly, monobody 12D5 stabilizes the H95/Y96 pocket, reducing its flexibility and altering the local environment to disrupt effector binding. We found that hydrophobic interactions play a central role in driving the binding affinity of KRAS complexes with monobodies and affimers. For instance, the hydrophobic cluster formed by W43 of affimer K6 with V7/I55/L56/M67 sites on KRAS is critical for binding. Similarly, affimer K3 stabilizes its interaction with KRAS through hydrophobic packing between KRAS hotspots (H95/Y96) and K3 hotspots (I41/I43/W44). These findings align with the general principle that hydrophobic interactions are critical for protein-protein interactions, especially in transiently exposed pockets like the SWII and H95/Y96 regions. The prominence of hydrophobic hotspots in these interactions suggests that small molecules or peptide-based inhibitors designed to mimic these interactions could achieve high binding specificity and affinity. Furthermore, the interplay between hydrophobic contacts and hydrogen bonding networks provides a blueprint for designing inhibitors that combine multiple interaction types to enhance stability and specificity.

The identification of key hotspot residues, such as H95 and Y96, may have useful implications for isoform-specific targeting. These residues, predominantly located in the α3 helix, are consistently implicated as major contributors to binding affinity and selectivity. For example, the unique presence of H95 in KRAS (and its absence in HRAS and NRAS) explains the isoform-specific recognition achieved by affimer K3. Mutational studies further validate the importance of H95, as substitutions mimicking HRAS/NRAS residues lead to significant destabilization and loss of binding. These findings highlight the potential of targeting isoform-specific residues to develop selective inhibitors for oncogenic KRAS mutants, addressing a longstanding challenge in cancer therapy. The agreement between computational predictions (mutational scanning and MM-GBSA analysis) and experimental data validates the robustness of these methodologies. Simplified energy models and free energy calculations accurately reproduce functional hotspots and predict binding energetics, providing a powerful framework for future studies. This convergence of computational and experimental approaches strengthens the reliability of the findings.

The allosteric network analysis of KRAS complexes with monobodies and affimer proteins provides critical insights into the structural and functional determinants of long-range communication in KRAS. We identified conserved allosteric hotspots and pathways that mediate signal transmission across functional regions of KRAS. Our analysis reveals a set of highly conserved residues that function as critical nodes in the allosteric communication network of KRAS. Residues in β-strand 4 (F78, L80, F82), α-helix 3 (I93, H95, Y96), β-strand 5 (V114, N116), and α-helix 5 (Y157, L159, R164) consistently emerge as hotspots across diverse binding partners. The predicted allosteric hotspots are in strong agreement with experimental data, which independently identified residues in β-strand 4 (F78, G79, F82) and β-strand 5 (V112, V114, N116) as critical mediators of long-range interactions affecting effector protein binding. This robust alignment between computational predictions and experimental findings underscores the reliability of the network-based approach in identifying functionally significant residues.

However, there many challenges remaining to fully understand mechanisms driving specific KRA binding and emergence of cryptic binding sites for allosteric intervention. First, the specificity of these interactions must be carefully optimized to avoid off-target effects. For instance, while the H95/Y96 pocket is unique to KRAS, subtle differences in amino acid composition between RAS isoforms necessitate precise targeting strategies. Second, the long-range effects induced by affimers like K69 raise questions about how such global conformational changes might influence other cellular processes. Addressing these challenges will require integrating computational modeling, structural biology, and experimental validation to refine inhibitor design. The ability of monobodies and affimers to induce localized perturbations that propagate globally across KRAS highlights the interconnected nature of protein structures. This interconnectivity suggests that even subtle changes in one region can have profound effects on distant functional sites, a principle that extends beyond KRAS to other signaling proteins. Moreover, the compensatory behavior observed in KRAS complexes—where stabilization of one region leads to increased flexibility in another—underscores the adaptability of proteins in response to external perturbations. Such insights could inform efforts to engineer synthetic binding partners or develop novel therapeutic strategies that exploit these dynamic properties.

## Conclusions

This study provides a comprehensive exploration of the dynamic, energetic, and allosteric mechanisms governing KRAS interactions with monobodies (12D1, 12D5) and affimer proteins (K6, K3, K69), offering critical insights into targeting this historically “undruggable” protein. Through an integrative approach combining molecular dynamics simulations, mutational scanning, binding free energy analysis, and network-based modeling, we identified conserved allosteric hotspots and pathways that mediate long-range communication across KRAS functional regions. The study reveals the intricate balance between structural rigidity and functional flexibility in KRAS, where stabilization of one region often induces compensatory mobility in others. For instance, monobody 12D1 rigidifies the SWII region, leading to enhanced SWI mobility, while affimer K3 leverages the α3 helix as a hinge point to amplify its effects. These observations emphasize the adaptability of KRAS and provide a blueprint for designing inhibitors that exploit transiently exposed cryptic pockets to stabilize inactive conformations. Importantly, the hydrophobic interactions driving these binding events offer a mechanistic foundation for developing small-molecule or peptide-based inhibitors with high specificity and affinity. The interplay between hydrophobic contacts and hydrogen bonding networks further highlights the multifaceted nature of KRAS interactions, suggesting that inhibitors combining multiple interaction types could achieve enhanced stability and selectivity. The network analysis of allostery reveals a set of highly conserved residues that function as critical nodes in the allosteric communication network of KRAS. A key outcome of this work is the identification of residues in β-strand 4 (F78, L80, F82), α-helix 3 (I93, H95, Y96), β-strand 5 (V114, N116), and α-helix 5 (Y157, L159, R164) as universal hotspots for both binding and allostery. The SWII/α3 pocket, particularly residues H95 and Y96, emerges as a pivotal control point for long-range communication, consistent with its role as a target for clinically approved inhibitors like sotorasib. This convergence of computational predictions with experimental evidence underscores the robustness of our methodologies and their potential to guide the discovery of isoform-specific inhibitors. The diversity of binding mechanisms and the identification of critical hotspots underscore the complexity of KRAS regulation and highlight the need for tailored approaches to drug discovery.

## Author Contributions

Conceptualization, G.V.; methodology, G.V.; software, M.A., V.P., B.F., G.H., G.V.; validation, G.V.; formal analysis, G.V., M.A., G.H., investigation, G.V.; resources, G.V., M.A. and G.H.; data curation, G.V.; writing—original draft preparation, G.V.; writing—review and editing, G.V., M.A. and G.H.; visualization, V.P., B.F., G.V.; supervision, G.V.; project administration, G.V.; funding acquisition, G.V. All authors have read and agreed to the published version of the manuscript.

## Conflicts of Interest

The authors declare no conflict of interest. The funders had no role in the design of the study; in the collection, analyses, or interpretation of data; in the writing of the manuscript; or in the decision to publish the results.

## Funding

This research was supported by the National Institutes of Health under Award 1R01AI181600-01 and Subaward 6069-SC24-11 and the Kay Family Foundation Grant A20-0032 to G.V.

## Data Availability Statement

Data is fully contained within the article and Supplementary Materials. Crystal structures were obtained and downloaded from the Protein Data Bank (http://www.rcsb.org). The rendering of protein structures was done with UCSF ChimeraX package (https://www.rbvi.ucsf.edu/chimerax/) and Pymol (https://pymol.org/2/).

## Acknowledgments

G.V acknowledges support from Schmid College of Science and Technology at Chapman University for providing computing resources at the Keck Center for Science and Engineering.

## Notes

### Competing Interest Statement

The authors have declared no competing interest.

