## Supplemental Figures S1-S5 for "Atomistic Profiling of KRAS Interactions with Monobodies and Affimer Proteins Through Ensemble-Based Mutational Scanning Unveils Conserved Residue Networks Linking Cryptic Pockets and Regulating Mechanisms of Binding, Specificity and Allostery"

**Supp****lementary Materials**

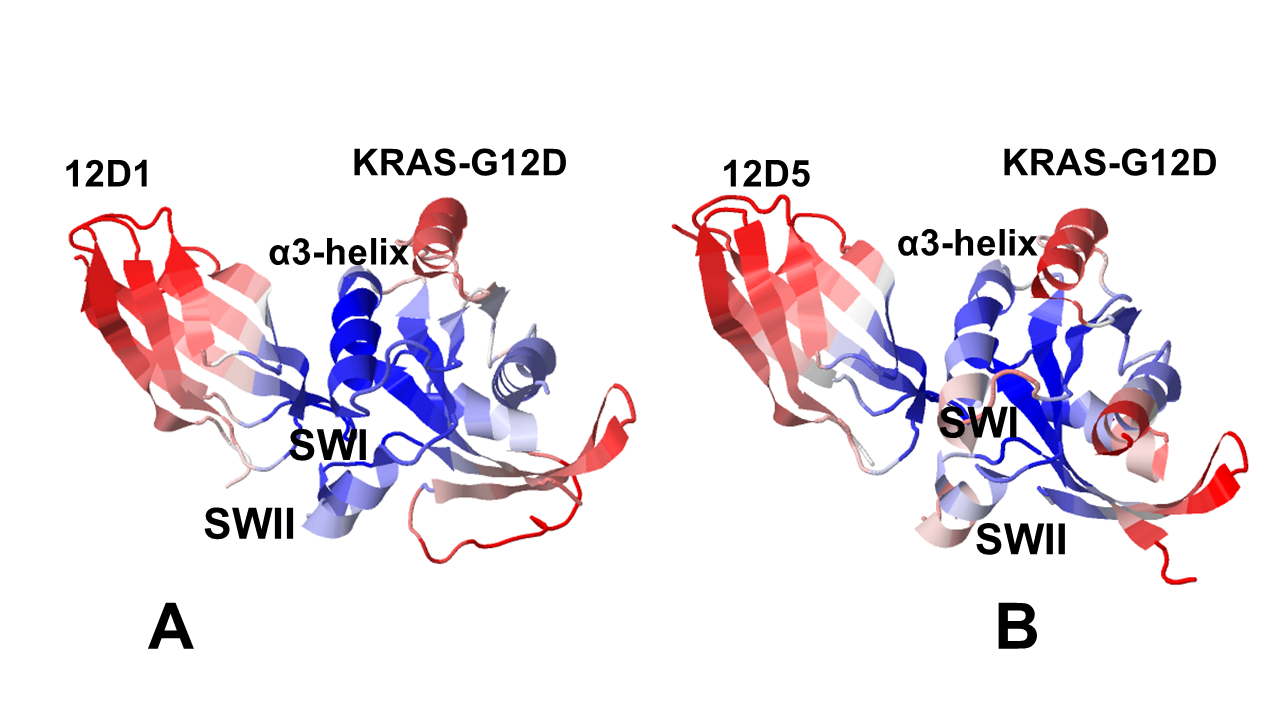

**Figure S1.** Structural maps of the essential mobility profiles for the KRAS-G1D complexes with monobody 12D1 (A) and monobody 12D5 (B). The essential mobility profiles are averaged over the first three major low frequency modes. The structures are shown in ribbons with the rigidity-to-flexibility scale colored from blue to red**.**

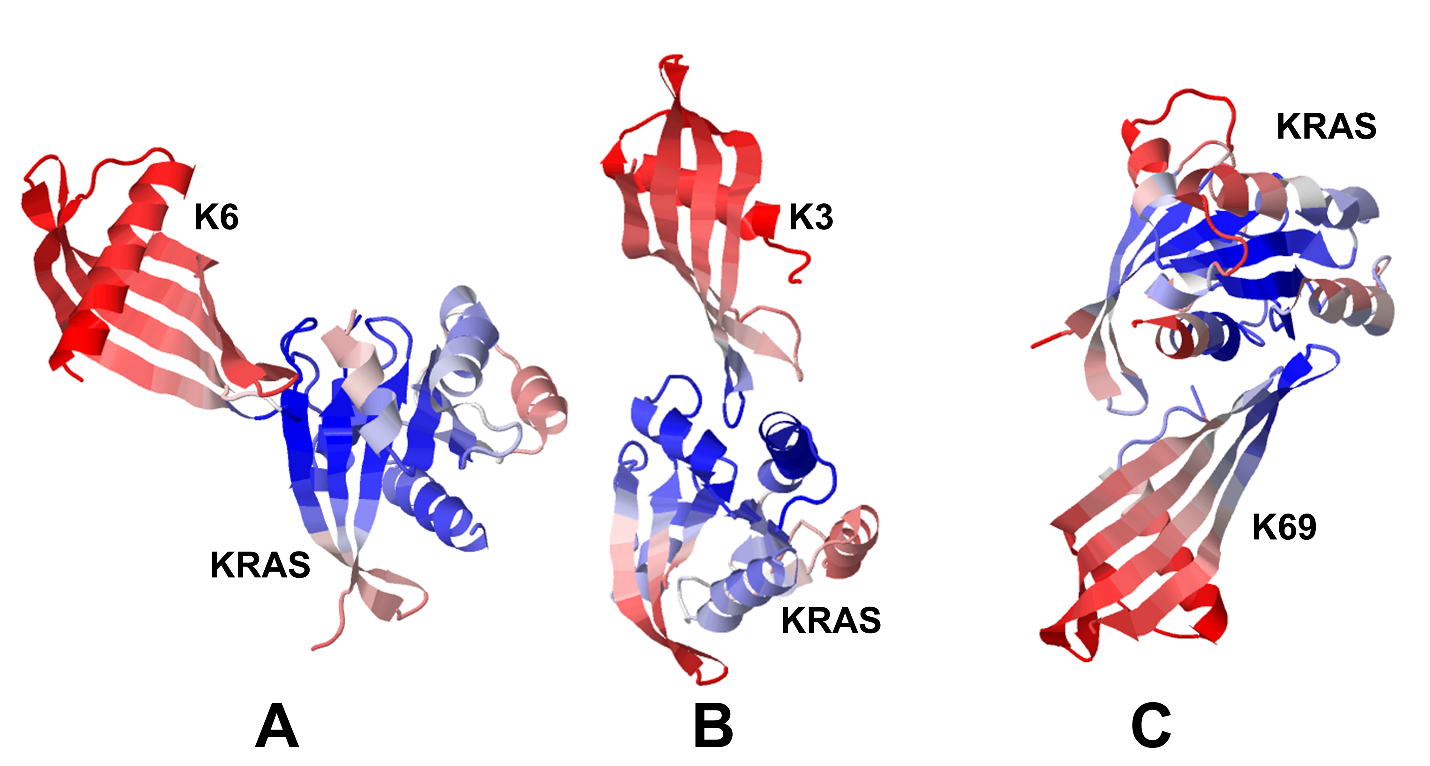

**Figure S2.** Structural maps of the essential mobility profiles for the KRAS-G1D complexes with affimer protein K6 (A), affimer protein K3 (B) and affimer protein K69 (C). The essential mobility profiles are averaged over the first three major low frequency modes. The structures are shown in ribbons with the rigidity-to-flexibility scale colored from blue to red**.**

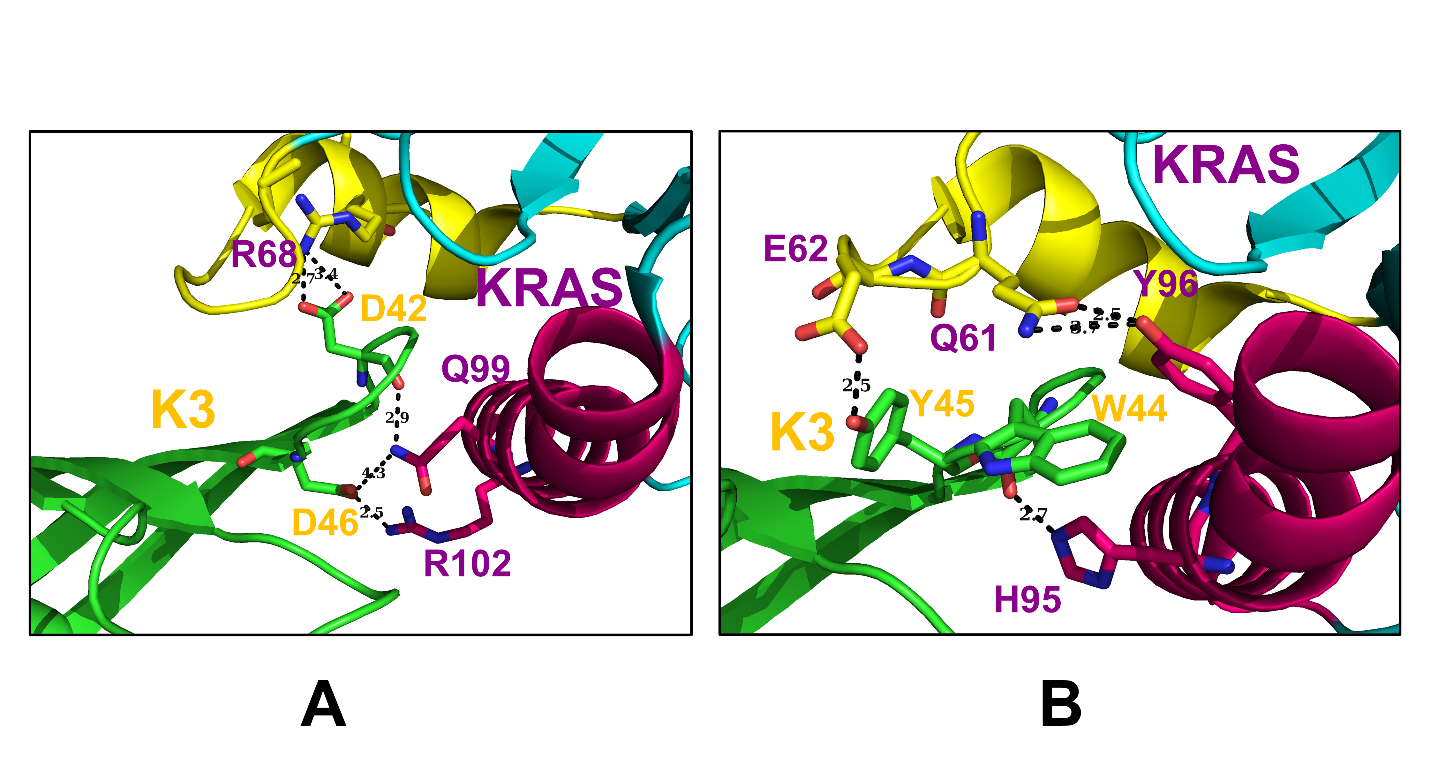

**Figure S3.** The interaction networks between KRAS and affimer K3 (A) Affimer K3 residues D42 and D46 (in orange ) bind to R68 of SII α- 3 (dark purple) and Q99 and R102 of α-3 (dark purple) bringing the two α helices in proximity. (B) The intermolecular interactions formed by W44 side chain and Y45 (in orange) of K3 burying into hydrophobic pocket and forming hydrogen bond with H95 of α-helix 3 (in dark purple) and E62 of SII (in dark purple).

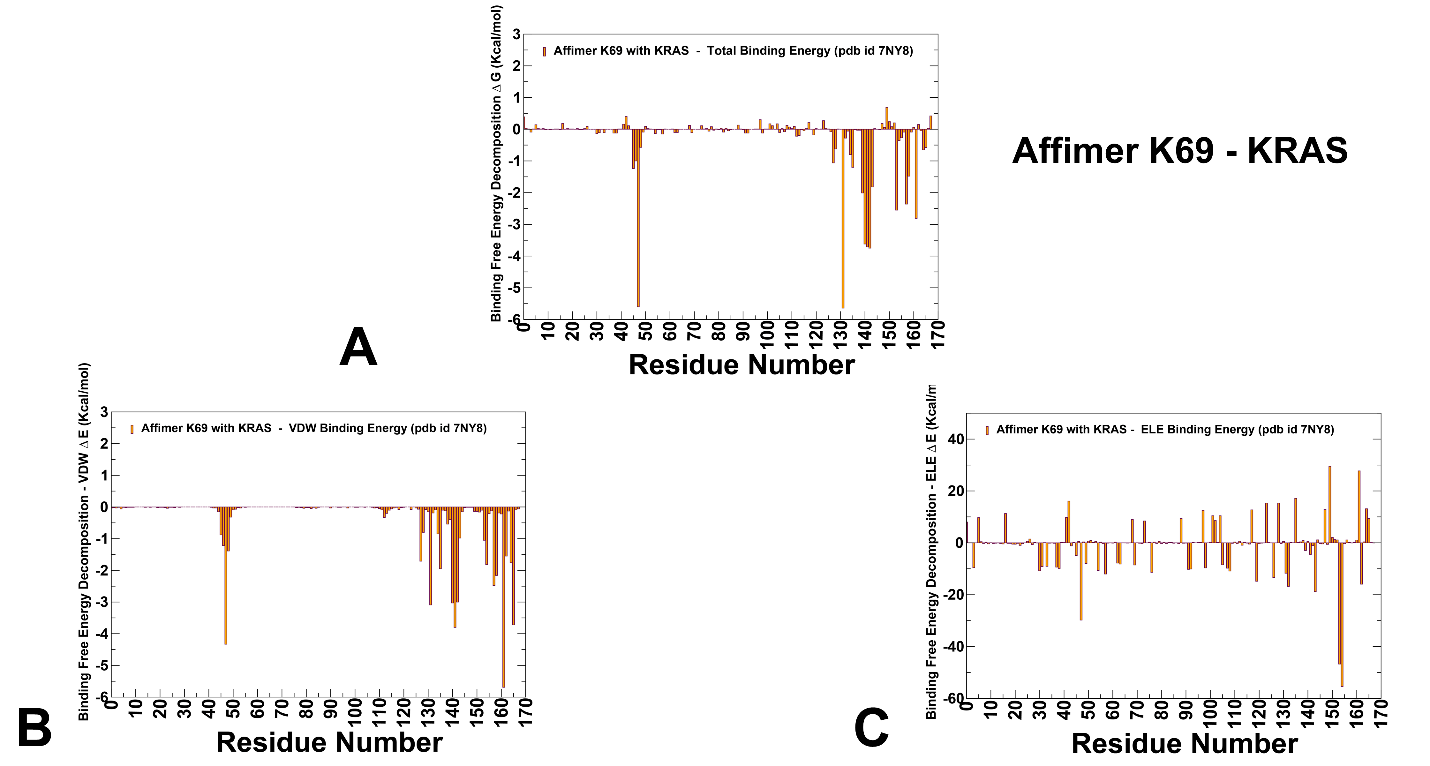

**Figure S4.** The residue-based decomposition of the total binding MM-GBSA energies for the KRAS residues in the KRAS complex with affimer K69. The residue-based decomposition of the total binding energy (A), van der Waals contribution (B) and electrostatic contribution to the total The MM-GBSA contributions are evaluated using 1,000 samples from the equilibrium MD simulations of KRAS-K69 complex.

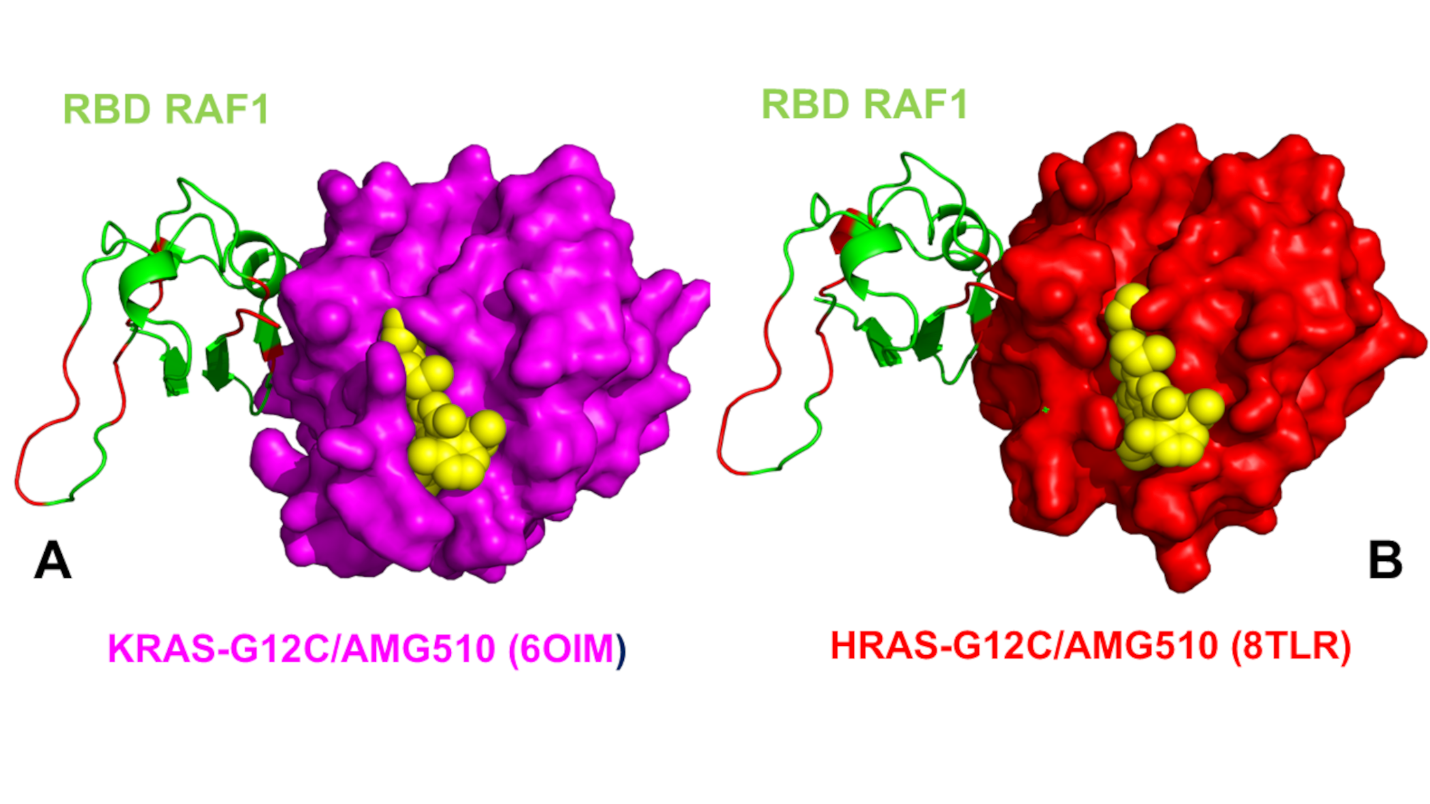

**Figure S5.** (A) The crystal structure of the human KRAS-G12C complex with RAF1 and allosteric inhibitor AMG510 KRAS-G12C is in magenta surface. RBD RAF1 is in green ribbons. AMG510 is in yellow spheres. The crystal structure of the human KRAS-G12C complex with RAF1 and allosteric inhibitor AMG510. (B) The crystal structure of the HRAS-G12C complex with RAF1 and allosteric inhibitor AMG510. HRAS-G12C complex (red surface), RBD RAF1 is in green ribbons and bound allosteric inhibitor AMG510 is yellow spheres.
